# SemiBin: Incorporating information from reference genomes with semi-supervised deep learning leads to better metagenomic assembled genomes (MAGs)

**DOI:** 10.1101/2021.08.16.456517

**Authors:** Shaojun Pan, Chengkai Zhu, Xing-Ming Zhao, Luis Pedro Coelho

## Abstract

Metagenomic binning is the step in building metagenome-assembled genomes (MAGs) when sequences predicted to originate from the same genome are automatically grouped together. The most widely-used methods for binning are reference-independent, operating *de novo* and allow the recovery of genomes from previously unsampled clades. However, they do not leverage the knowledge in existing databases. Here, we propose SemiBin, an open source tool that uses neural networks to implement a semi-supervised approach, *i.e.* SemiBin exploits the information in reference genomes, while retaining the capability of binning genomes that are outside the reference dataset. SemiBin outperforms existing state-of-the-art binning methods in simulated and real microbiome datasets across three different environments (human gut, dog gut, and marine microbiomes). SemiBin returns more high-quality bins with larger taxonomic diversity, including more distinct genera and species. SemiBin is available as open source software at https://github.com/BigDataBiology/SemiBin/.

## 1 Main

DNA sequences obtained from mixed communities (the metagenome) can be computationally grouped into *bins*, such that each bin is predicted to contain only sequences from the same genome. This process, known as binning, has enabled the construction of large compendia of metagenome-assembled genomes (MAGs) from human-associated (Almeida et al., 2019, 2020; Pasolli et al., 2019), animal-associated (Stewart et al., 2019), and environmental (Tully et al., 2018) samples.

Binning methods can be divided into two categories, depending on whether they rely on pre-existing genomes (reference-dependent) or operate *de novo* (reference-independent). Reference-dependent methods are limited to discovering novel genomes from previously-known species, while reference-independent ones can recover completely novel species and even novel phyla (Parks et al., 2017; Nascimento Lemos et al., 2020). Most reference-independent methods are completely unsupervised as they do not use information from reference genomes, instead relying on sequence features (*e.g. k*-mer frequencies) and abundance, which are assumed to vary more between than within genomes. One exception is SolidBin (Wang et al., 2019), which uses a semi-supervised approach (Gu et al., 2012). In particular, SolidBin takes advantage of *must-link* and *cannot-link* constraints between pairs of contigs: a *must-link* constraint indicates that the two contigs should be binned together, while a *cannot-link* one indicates that they should not (despite their name, these constraints are not strictly followed by the algorithm). SolidBin generates these constraints by alignment to existing genomes, marking pairs of contigs that align to the same species as *must-link*, while pairs that align to different genera are considered as *cannot-link*.

Here, we introduce a semi-supervised binning method, SemiBin (Semi-supervised metagenomic Binning), based on deep contrastive learning to take advantage of *must-link* and *cannot-link* constraints (Śmieja et al., 2020) (see Fig. 1a and Supplementary Fig. 1). Generating these constraints by alignment to reference genomes is subject to both noise (from annotation error) and sampling bias in genome databases (see Supplementary Text). In particular, while *cannot-link* constraints could be robustly derived from contig annotations to the GTDB (Parks et al., 2020), *must-link* constraints are instead generated by breaking up longer contigs artificially (see Supplementary Text). The constraints serve as input to a semi-supervised siamese neural network (Śmieja et al., 2020) for feature embedding. SemiBin then generates a sparse graph from the embedded features and groups the contigs into bins with the Infomap (Rosvall and Bergstrom, 2008) community detection algorithm (see Methods).

**Fig 1.**
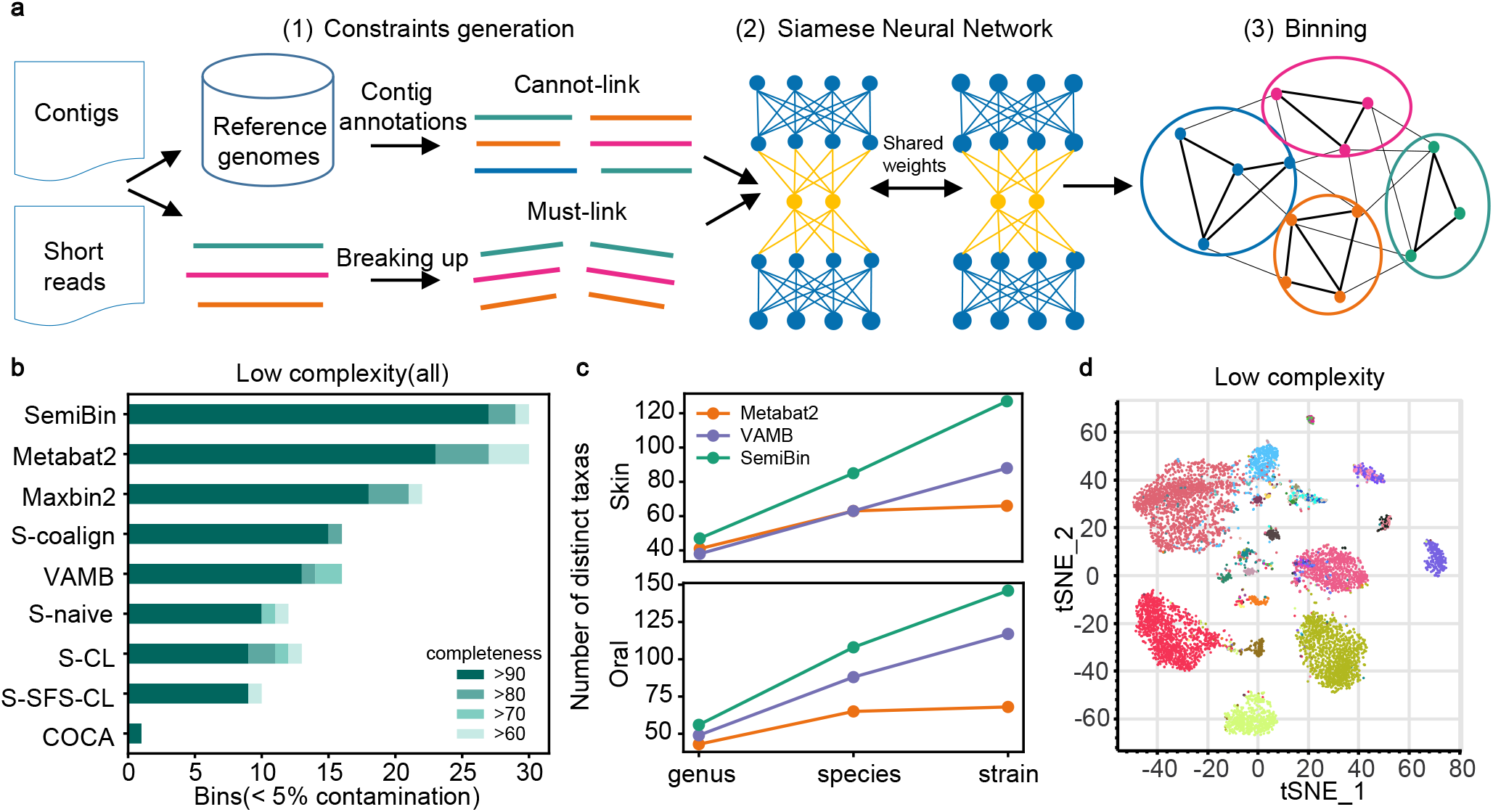
SemiBin outperformed other binners in simulated datasets with single-sample, co-assembly and multi-sample binning. **a,** An overview of SemiBin. (1) *Must-link* constraints are generated by artificially breaking up contigs, and *cannot-link* constraints are generated by contig taxonomic annotations. (2) These constraints are used as input to a semi-supervised deep learning model to learn an embedding (see Supplementary Fig. 3). (3) Contigs are binned on the graph created by considering contigs as vertices with edge weights derived from the embedding learned (see Methods). **b,** In CAMI I simulated datasets, SemiBin returned more high-quality bins. Shown is the number of reconstructed genomes per method with varying completeness and contamination < 5% (methods shown are, top to bottom: SemiBin, Metabat2, Maxbin2, SolidBin-coalign, VAMB, SolidBin-naive, SolidBin-CL, SolidBin-SFS-CL, and COCACOLA). SemiBin also outperforms all other methods in the medium and high complexity datasets (see Supplementary Fig. 5a). **c,** SemiBin reconstructed a larger number of distinct high-quality genera, species and strains in the CAMI II Skin and Oral datasets compared to either Metabat2 or VAMB. A high-quality strain is considered to have been reconstructed if any bin contains the strain with completeness > 90% and contamination < 5% (see Methods). If at least one high-quality strain is reconstructed for a particular genus or species, then those are considered to have been reconstructed. **d,** Semi-supervised embedding separates contigs from different genomes. Shown is a two-dimensional visualization of embedding of the low complexity dataset from CAMI I, with contigs colored by their original genome (using t-SNE, we used Sklearn.manifold.TSNE(perplexity=50,init = pca)) (Pedregosa et al., 2011)

As studies often collect multiple related metagenomes, there have been different proposals on how to handle binning in this context: *single-sample* (where each metagenome is handled independently), *co-assembly* (where different metagenomes are pooled together), or *multi-sample* (where resulting bins are sample-specific, but abundance information is aggregated across samples) (Nissen et al.). All three modes are supported by SemiBin.

To compare the performance of SemiBin to existing binners, we first benchmarked SemiBin, Metabat2 (Kang et al., 2019), Maxbin2 (Wu et al., 2016), VAMB (Nissen et al.), SolidBin (Wang et al., 2019), and COCACOLA (Lu et al., 2017) on the five simulated datasets from CAMI I and CAMI II (Sczyrba et al., 2017; Meyer et al., 2021) (the Critical Assessment of Metagenome Interpretation). The CAMI I datasets comprise different numbers of organisms including strain variation with low (40 genomes, 1 sample), medium (132 genomes, 2 samples), and high (596 genomes, 5 samples) complexity and were used to evaluate single-sample (low complexity) and co-assembly (medium and high complexity) binning. CAMI II datasets mimic different human body sites and contain multiple simulated metagenomes from the same environments, thus allowing us to test multi-sample binning.

In the CAMI I datasets, SemiBin was able to reconstruct the highest number of high-quality bins (completeness > 90% and contamination < 5% (Bowers et al., 2017)) and achieved the best F1-score (see Fig. 1b, Supplementary Figs. 4 and 5). SemiBin reconstructed 17.4%, 11.4% and 11.4% more high-quality bins compared to the second highest result in the low, medium, and high complexity datasets respectively. Since metagenomic binning is particularly challenging when multiple strains from the same species are present (Brown et al., 2020), we evaluated the performance of the methods on common strains (defined as genomes for which another genome with at least 95% average nucleotide identity (ANI) is present in the dataset (Sczyrba et al., 2017)) in the three datasets. SemiBin was able to reconstruct 11.1%, 18.5% and 35.3% more high-quality common strains than the second-best alternative (see Supplementary Fig. 5).

In the CAMI II datasets, we compared SemiBin to VAMB and Metabat2 on the Skin (610 genomes, 10 samples) and Oral (799 genomes, 10 samples) datasets, as previous benchmarking studies had indicated their best performance on these (Nissen et al.). We benchmarked the original Metabat2 and an adapted version which implements multi-sample binning (see Methods). This adaptation, however, led only to modest improvements (see Supplementary Fig. 8a). Compared to VAMB (the second best binner), SemiBin could reconstruct 44.3% and 24.8% more distinct high-quality strains, 34.9% and 22.7% more distinct species and 23.7% and 14.3% more distinct genera in the Skin and Oral datasets respectively (see Fig. 1c). We evaluated the behavior of the methods on multi-strain species and observed that SemiBin could reconstruct more high-quality distinct strains across almost all ANI intervals, even when very similar genomes (ANI > 99.5%) are present (see Supplementary Fig. 9).

To show the impact of semi-supervised deep learning, we compared the proposed method to the same pipeline without the deep learning feature embedding step (see Methods). SemiBin could reconstruct 6.7%-65.0% more distinct, high-quality bins compared to this non-embedding version in the five datasets, and the improvement was especially pronounced in complex environments (see Supplementary Fig. 10 and 12). SemiBin with semi-supervised learning also outperformed a modified version of SemiBin that directly used the *must-link* and *cannot-link* constraints to generate the graph for binning (see Supplementary Fig. 11, 12, Methods and Supplementary text). These results showed that the semi-supervised learning step, besides remembering the inputs, had the ability to learn the underlying structure of the environment from the *must-link* and *cannot-link* constraints.

To test SemiBin on real data, we applied it to datasets from three different environments: a human gut (n = 82) (Wirbel et al., 2019), a non-human animal-associated (dog gut, n = 129) (Coelho et al., 2018), and an environmental (ocean surface samples from the Tara Oceans project, n = 109) microbiome dataset (Sunagawa et al., 2015). In large-scale metagenomic analyses, single-sample binning is widely used (see Supplementary Table 3) because samples can be trivially processed in parallel. Here, we compared SemiBin to Maxbin2, VAMB and Metabat2 with single-sample binning. Additionally, because Nissen et al. (Nissen et al.) have demonstrated the effectiveness of multi-sample binning in real projects, we compared SemiBin with VAMB in this mode.

Because the actual genomes are unknown in real data, we relied on CheckM (Parks et al., 2015) and GUNC (Orakov et al., 2021) to evaluate the quality of the recovered bins. In all cases, SemiBin recovered more high-quality bins than the alternatives considered (see Fig. 2b and Supplementary Fig. 14). In the human gut, dog gut, and ocean datasets, SemiBin reconstructed 1497, 2415 and 446 high-quality bins with single-sample binning, significantly outperforming Metabat2 with an increase of 437 (41.2%), 1011 (72.0%) and 146 (48.7%), respectively. For multi-sample binning, SemiBin could reconstruct 17.5%, 11.0% and 30.7% more high-quality bins than VAMB. In the human gut and ocean datasets, SemiBin with single-sample binning performed better than VAMB with multi-sample binning, which requires more computational time for short-read mapping and cannot be performed in parallel. We annotated these high-quality bins from multi-sample binning with GTDB-Tk (Chaumeil et al., 2020). SemiBin could reconstruct more distinct taxa when compared to VAMB in the human gut and ocean datasets, performing similarly in the dog gut dataset (see Supplementary Fig. 16).

**Fig 2.**
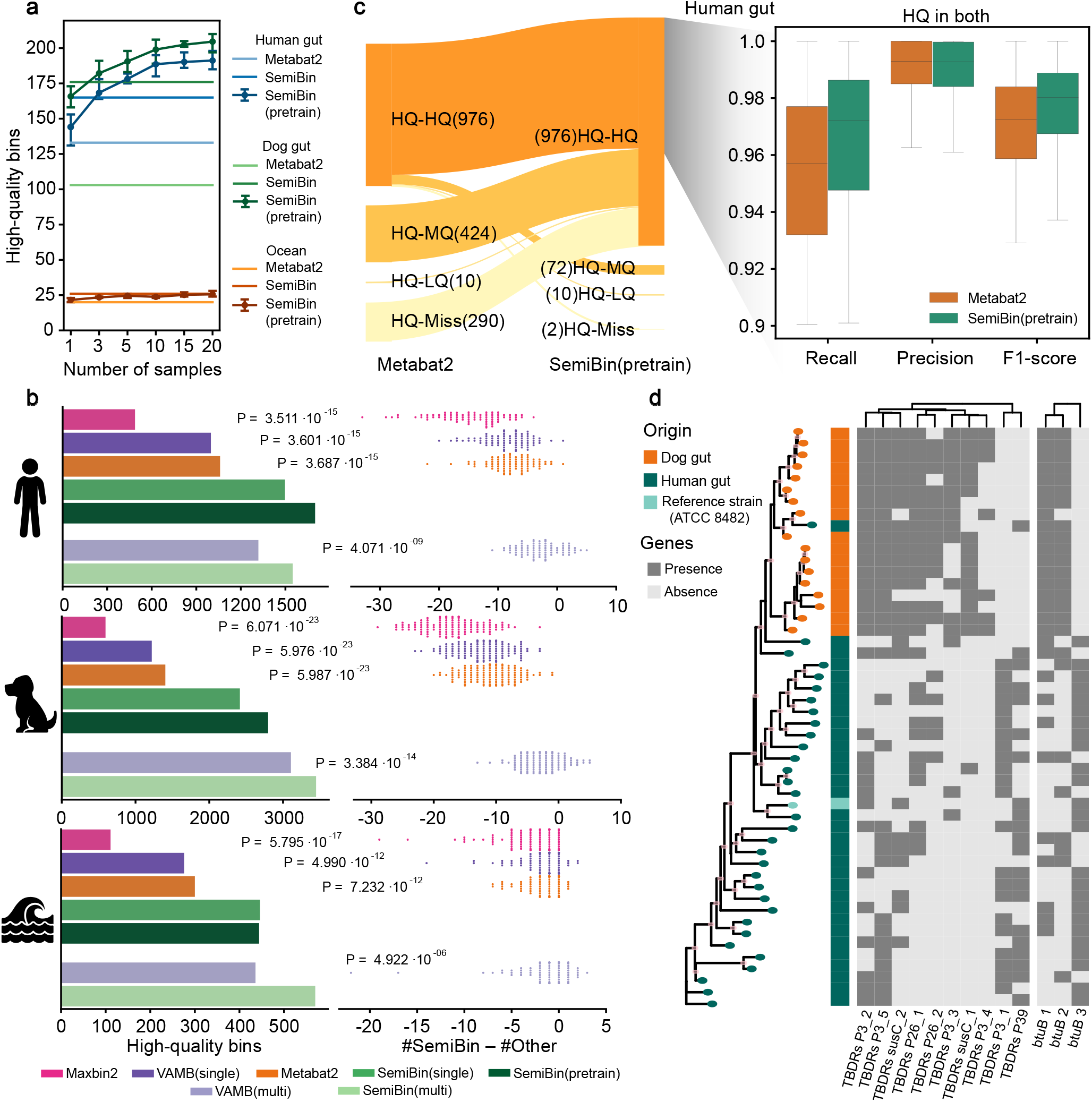
SemiBin outperformed other binners in real datasets. **a,**Number of high-quality bins (automatically evaluated using CheckM and GUNC, see Methods) using a pretrained model with different numbers of samples used for training. Per-sample SemiBin and Metabat2 also shown for comparison. The average, worst and best result from five runs are shown for the pretrained model. **b,** SemiBin(pretrain) produced more high-quality bins compared to Maxbin2, VAMB and Metabat2 in the human gut, the dog gut, and the ocean datasets (**left**). When considering the results in each sample (**right**), SemiBin(pretrain) outperformed other single-sample binners in almost every sample. In the dog gut dataset, SemiBin(pretrain) always produced more high-quality bins than any other per-sample methods. In the human gut and ocean datasets, other methods occasionally outperformed SemiBin(pretrain) (observed for 2 human gut samples and 9 ocean samples), but the difference is never large (at most, two extra high-quality bins were produced). Results of VAMB in multi-sample binning mode are compared to SemiBin(multi), with SemiBin(multi) producing more high-quality bins overall (although not in every sample). P-values shown are from a Wilcoxon signed-rank test (two-sided) on the counts for each sample. **c,** We identified the overlap between the bins in the human gut from Metabat2 and SemiBin(pretrain) using Mash (see Methods). While some high-quality bins from Metabat2 were matched to lower quality SemiBin(pretrain) bins, the reverse was much more common (i.e. SemiBin(pretrain) recovered higher-quality versions of bins that were only medium-quality in the Metabat2 outputs as well as some that were completely absent). Within bins that are present at high-quality in the output of both binners, SemiBin(pretrain) achieved higher completeness (recall) (*P* = 2.236ů10^−67^) without a statistically-significant increase in contamination (1 - precision) (*P* = 0.167). The overall F1 statistic (2Œ(recall Œ precision)/(recall + precision)) is significantly better (*P* = 9.287ů10^−67^; all P-values were computed from using Wilcoxon signed rank test, two-sided null hypothesis). Results in the dog gut and ocean datasets are qualitatively similar (see Supplementary Fig. 15). (HQ-HQ: high-quality in both; HQ-MQ: high-quality in one and medium-quality in the other; HQ-LQ: high-quality in one and low-quality or worse in the other; HQ-Miss: high-quality in one and missed in the other) **d,** Strains of *B. vulgatus* recovered from the human and dog gut microbiomes and a type strain *B. vulgatus* ATCC 8482 cluster according to their host. Shown is the maximum-likelihood phylogenetic tree based on core genes; branches with bootstrap values higher than 70 were marked pink in the nodes. Clustering based on ANI or gene presence also showed a separation between the hosts (see Supplementary Fig. 21). The gene content of the strains was also statistically different between the two hosts (*P* < 0.05 using Fishers exact test after FDR correction using Benjamini-Hochberg method, see Methods), in particular in the presence of different TonB-dependent receptors (TBDR) and their interaction partner *btuB*, which are involved in nutrient uptake (see Supplementary Text)

Generating the cannot-link constraints and training the models comes at a substantial computational cost (see Supplementary Table 4) and, by default, SemiBin learns a new embedding model for each sample. Therefore, for single-sample binning, the possibility of reusing a learned model from one sample to another (*model transfer*) was tested with encouraging results (see Supplementary Fig. 13 and Supplementary text). However, there was still a loss in the number of high-quality bins recovered compared to learning a new model for every sample. To overcome this, we built models from multiple samples and verified that this could obtain the best performance, with the scalability of low computational per-sample costs (see Fig. 2a and Supplementary. Table 4), an approach we termed SemiBin(pretrain).

If a sufficient number of samples was used in pretraining (see Methods), SemiBin(pretrain) could reconstruct more high-quality bins, and SemiBin(pretrain) outperformed Metabat2 in all cases (see Fig. 2a). In the human gut and dog gut datasets, SemiBin(pretrain) trained on more than three samples outperformed the original SemiBin and performed at a similar level in the ocean dataset. As pretraining is performed only once and applying the model to other samples is computationally fast, SemiBin(pretrain) can be used in large-scale metagenomic analysis (see Supplementary Table 4).

For every environment, we also benchmarked SemiBin(pretrain) with the best performing pretrained model (see Fig. 2b). Compared to the original SemBin, SemiBin(pretrain) could reconstruct 203 (13.6%) and 382 (15.8%) more high-quality bins in human and dog gut datasets and achieved similar results in the ocean dataset. When compared to Metabat2, SemiBin(pretrain) could reconstruct 640 (60.4%), 1393 (99.2%) and 144 (48.0%) more high-quality bins. SemiBin (pretrain) also performed significantly better than Metabat2 when comparing high-quality bins on a sample-by-sample basis (Wilcoxon signed-rank test, two-sided, *P* = 3.687ů10^−15^ (n = 82), *P* = 5.987ů10^−23^ (n = 129), *P* = 7.232ů10^−12^ (n = 109); see Fig. 2b).

We used Mash (Ondov et al., 2016) to identify instances when SemiBin(pretrain) and Metabat2 reconstructed the same genome. Most Metabat2-generated high-quality bins corresponded to high-quality bins generated by SemiBin(pretrain). SemiBin(pretrain) results further contained many high-quality bins which corresponded to lower-quality (or absent) bins in the Metabat2 results, the inverse case (Metabat2 generating a higher-quality version of a SemiBin(pretrain) bin) being relatively rare. For genomes that were recovered at high-quality by both binners, the SemiBin(pretrain) bins had, on average, higher completeness and F1-score, with only a minimal increase in contamination that was not statistically significant in the human and dog gut datasets (see Fig. 2c and Supplementary Fig. 15).

We annotated the high-quality bins generated by SemiBin(pretrain) and Metabat2 with GTDB-Tk (Chaumeil et al., 2020). Bins from SemiBin(pretrain) represented a larger taxonomic diversity at all levels (see Supplementary Fig. 17). Additionally, SemiBin(pretrain) was able to recover more genomes from both known and unknown species (see Supplementary Fig. 18), validating the ability of the semi-supervised embedding to extract information from the background data while being capable of going beyond it.

To further test the generalization ability of the learned model, we applied the best-performing pretrained model in the human gut dataset (termed as SemiBin(pretrain; external), see Methods) to two human gut datasets not used for model training (hold-out datasets). As the human gut dataset considered so far is from a German population originated sample (where individuals are assumed to consume a Western-style diet), we tested both on another dataset from the German population (Louis et al., 2016) and on a dataset from a non-Westernized African human population (Pasolli et al., 2019). All versions of SemiBin (see Methods) achieved excellent results compared to Metabat2 and SemiBin (pretrain; external) significantly outperformed Metabat2 (Wilcoxon signed-rank test, two-sided, *P* = 3.050ů10^−12^ (n = 92), *P* = 1.014ů10^−7^ (n = 50); see Supplementary Fig. 19), showing that the pretrained model could be used on hold-out datasets.

Amongst species shared between the human and dog gut microbiomes in our study, *Bacteroides vulgatus* had the largest number of recovered strains in both. Namely, 17 dog gut and 32 human gut *B. vulgatus* bins were obtained with SemiBin(pretrain) (compared to four and five obtained with Metabat2). Based on either sequence similarity or gene content, the bins from the dog gut clustered separately from those of the human gut (see Fig. 2d, Supplementary Fig. 21). In particular, we found different gene presence patterns for genes encoding TonB-dependent receptors(TBDR), ten of which showed statistically significant differences (after multiple hypothesis correction; see Fig. 2d, Supplementary Table. 5). We also found that the three *btuB* genes in dog and human gut microbiomes (see Fig. 2d and Supplementary Table. 5) encoding a protein that interacts with TonB (Chimento et al., 2003) belong to different orthologous groups. As different btuB-TonB complexes may affect the transport of essential micronutrients such as vitamin B12 (Shultis et al., 2006), *Bacteroides vulgatus* in human and dog gut microbiomes might have different degrees of ability to transport vitamin B12.

We showed the superior results of SemiBin when compared to other state-of-the-art binning methods, demonstrating the advantages of using background knowledge in metagenomic binning. Looking forward, we expect that genome databases will continue to improve in quality along with taxonomic prediction methods. This should make the use of this information even more valuable and further widen the gap with respect to purely *de novo* methods.

## 2 Methods

### 2.1 SemiBin pipeline

The input to SemiBin consists of a set of contigs and mapped short reads (in the form of BAM files) from one or multiple samples. The output consists of a collection of bins with the original contigs clustered.

#### 2.1.1 Preprocessing

Prior to binning, contigs are filtered to remove short contigs, with the default threshold being 2,500bp. If the contigs in the range 1,0002,500bp represent < 5% of the total number of basepairs (this parameter can be changed by the user), then the threshold is reduced to 1,000bp.

After filtering, every contig is represented by *k*-mer relative frequencies *K* and abundance distribution *A*. SemiBin uses *k* = 4 (also used in other tools (Wu et al., 2016; Kang et al., 2019; Nissen et al.; Lu et al., 2017; Wang et al., 2019)), so the dimension of *K* is 136 (*k*-mers that are the reverse complement of each other are considered equivalent). *K*-mer vectors are normalized by the lengths of each contig. We defined:

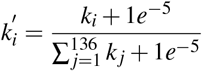

The abundance *a* is defined as the average number of reads per base mapping to the contig. If the number of samples *N* used in the binning is greater than or equal to 5, SemiBin scales the abundance features from the *N* samples to a similar order of magnitude as the *k*-mer (which are restricted to the 0-1 range, by construction). In particular, the constant *s* used to scale was defined as 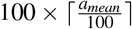. After the processing, the input to the semi-supervised model is the vector 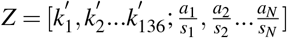. Otherwise, when the number of samples used is smaller than 5, the input to the semi-supervised model is the vector *Z* = [*k*′_1_, *k*′_2_ …*k*′_136_].

#### 2.1.2 Generating must-link and cannot-link constraints

SemiBin uses taxonomic annotation results to generate *cannot-link* constraints and break up contigs to build textitmust-link constraints. By default, MMseqs2 (Steinegger and Söding, 2017; Mirdita et al., 2020) (version 13.45111, default parameters) is used to annotate contigs with GTDB (Parks et al., 2020) reference genomes. We defined contigs pairs with different annotations at species level (with scores both above 0.95) or at genus level (with scores both above 0.80) as containing a *cannot-link* constraint between them. If the number of *cannot-link* constraints is above a threshold (default is 4 million), they are randomly sampled to speed up the training process. SemiBin also employs the same method as in Maxbin2 (Wu et al., 2016) to predict seed contigs for different species using single-copy marker genes and generate *cannot-link* constraints between them.

To generate *must-link* constraints, SemiBin breaks up long contigs into two fragments with equal length artificially and generates *must-link* constraints between the two shorter contigs. By default, SemiBin uses a heuristic method to automatically find the minimum size of contigs to break up (alternatively, the user can specify the threshold): it breaks up contigs that are at least as long as the longest contig such that the basepairs contained in all contigs that are as long (or longer) encompass ≥ 98% of the total number of basepairs (with an additional minimum size of 4,000bps).

#### 2.1.3 Semi-supervised Siamese neural network architecture

We used a semi-supervised siamese neural network (see Supplementary Figs. 2 and 3) (Śmieja et al., 2020) dealing with the *must-link* and *cannot-link* constraints. The network has two siamese neural networks with shared weights.

As it is semi-supervised, this neural network is trained with two loss functions. The supervised loss is a contrastive loss which is used to classify pairs of inputs as *must-link* or *cannot-link*:

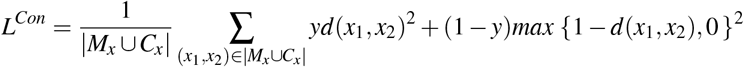

where *M_x_* denotes the *must-link* pairs and *C_x_* the *cannot-link* pairs in the training set, *d*(*x*_1_, *x*_2_) is the Euclidean distance of the embedding of (*x*_1_, *x*_2_), and *y* is an indicator variable for the (*x*_1_, *x*_2_) pair (with the value 1 if (*x*_1_, *x*_2_) ∈ *M_x_*, and 0 if (*x*_1_, *x*_2_) ∈ *C_x_*).

The goal of the supervised embedding is to transform the input space so that contigs have a smaller distance if they are in the same genome, compared to pairs of contigs from different genomes. To ensure that the embedding learns structure shared by all genomes, we also used an autoencoder (Hinton and Zemel, 1994) to reconstruct the original input from the embedding representation with an unsupervised mean square error (MSE) loss function:

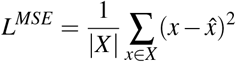

where *x* is the original input and 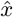 is the reconstructed input, *X* are whole contigs in the dataset.

To avoid overfitting, we used Dropout (Srivastava et al., 2014) and batch normalization (Ioffe and Szegedy, 2015) in the model and we also used leaky rectified linear unit (Maas et al., 2013) activation function.

The model used here is a dense deep neural network (see Supplementary Fig. 3). The dimension of the input features, *F* depends on the number of samples used in the binning (see “Preprocessing” Section). The first two layers of the encoding and the decoding network are followed by batch normalization, a leaky rectified linear unit and a dropout layer (the dropout rate used was 0.2). The purpose of the training is to optimize the contrastive loss and the MSE loss at the same time with Adam (Kingma and Ba, 2014) optimization algorithm. The model was implemented in Pytorch (Paszke et al., 2019).

#### 2.1.4 Similarity between contigs

The similarity between two contigs is defined as

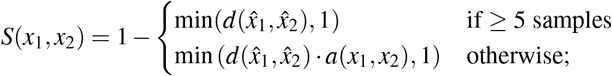

where 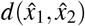 is the Euclidean distance of the semi-supervised embedding. When there are fewer than 5 samples, the embedding distance only contains *k*-mer information. In this case, we modeled the number of reads per base of one contig as a normal distribution, and used the Kullback-Leibler divergence (Kullback and Leibler, 1951) to measure the divergence between the normal distributions from contigs, denoted as *a*(*x*_1_, *x*_2_) above.

#### 2.1.5 Clustering

The binning problem is modeled as clustering on a graph. First, SemiBin considers the fully connected graph with contigs as nodes and *similarity* between contigs as the weight of edges. To convert the community detection task to an easier task, the fully connected graph is converted to a sparse graph. A parameter (*max_edges*, defaulting to 200) is used to control the sparsity of the graph, but the results are robust to different values of this parameter (see Supplementary Fig. 6 and 10). For each node, only the *max_edges* edges with the highest weights are kept. To remove any potential artefacts introduced by the embedding procedure, SemiBin builds another graph with the same procedure using the original features and the edges in the graph built from embedding that do not exist in the graph from original features are also removed.

After building the sparse graph, Infomap, an algorithm to reveal community structure in weighted graph based on information theory (Rosvall and Bergstrom, 2008), is used to reconstruct bins from the graph. If the user requests it, SemiBin can use single-copy genes of the reconstructed bins to independently re-bin bins whose mean number of single copy genes is greater than one (Wu et al., 2016). For this, SemiBin uses the weighted *k*-means algorithm to recluster bins according to the semi-supervised embedding and the abundance features. The results in the manuscript were obtained with this reclustering step. Finally, SemiBin outputs the final binning results, removing bins smaller than a user-definable threshold (which defaults to 200kbps).

#### 2.1.6 SemiBin(pretrain)

The default pipeline of SemiBin is to (1) generate *must-link*/*cannot-link* constraints, (2) train the semi-supervised deep learning model for this sample, (3) bin based on the embeddings. To address the issue that contig annotations and model training requires significant computational time and considering the *k*-mer frequencies can be transferred between samples or even project, we proposed SemiBin(pretrain) for single-sample binning that (1) we trained a model with constraints from one sample or several samples and (2) we directly applied this model to other samples. To use SemiBin(pretrain) in the tool, users can train a model from their datasets or use one of our built-in pretrained models for human gut, dot gut, and ocean environments.

#### 2.1.7 Binning modes

We have evaluated our model in three binning modes: single-sample, co-assembly, and multi-sample binning (Nissen et al.). Single-sample binning means binning each sample into inferred genomes after independent assembly. This mode allows for parallel binning of samples, but it does not use information across samples.

Co-assembly binning means samples are co-assembled first and then binning contigs with abundance information across samples. This mode can generate longer contigs and use co-abundance information, but co-assembly may lead to inter-sample chimeric contigs (Sangwan et al., 2016) and binning based on co-assembly cannot retain sample specific variation (Nissen et al.).

Multi-sample binning means the resulting bins are sample-specific (as in single-sample binning), but information is aggregated across samples (in our case, abundance information). This mode requires more computation resources as it requires mapping reads back to the concatenated FASTA file.

In single-sample and co-assembly binning, we calculate the *k*-mer frequencies and abundance for every sample and bin contigs from every sample independently. For multi-sample binning, we first concatenate contigs from every sample into a single database to which short reads are mapped to generate abundance profiles. Unlike what is used in VAMB (which introduced the *multi-sample* binning concept), this concatenated database is then re-split and contigs from each sample are binned separately.

### 2.2 Data used

For the benchmarking of binners, we used 5 simulated datasets from CAMI I and CAMI II and 5 real metagenomic datasets. Five simulated datasets of CAMI I and CAMI II were downloaded from CAMI challenge (Sczyrba et al., 2017; Meyer et al., 2021). CAMI I includes three datasets: low complexity, medium complexity, and high complexity datasets. The low complexity dataset has 40 genomes with a single simulated sample. Medium complexity dataset has 132 genomes with two simulated samples with different insert sizes. Here we used samples with 150bp insert size. The high complexity dataset has 596 genomes with 5 samples. We also used skin and oral cavity datasets in toy human microbiome project dataset of CAMI Skin dataset has 610 genomes with 10 samples and Oral dataset has 799 genomes with 10 samples. We used low complexity dataset to evaluate the single-sample binning mode of our method, medium and high complexity dataset to evaluate the co-assembly binning mode and Skin, Oral datasets to evaluate the multi-sample binning mode. We used fastANI (Jain et al., 2018) (version 1.32, default parameters) to calculate the ANI value between genomes for every sample from CAMI II datasets.

We also used five real microbiome projects from different environments to evaluate the proposed method:

1. a human gut dataset with 82 samples (Wirbel et al., 2019) (study accession PRJEB27928),
2. a dog gut dataset with 129 samples (Coelho et al., 2018) (study accession PRJEB20308),
3. a marine data from the Tara Oceans project with 109 ocean surface samples (Sunagawa et al., 2015) (study accession PRJEB1787),
4. a German human gut dataset with 92 samples (Louis et al., 2016) (study accession PRJNA290729),
5. a non-Westernized African human gut dataset with 50 samples (Pasolli et al., 2019) (study accession PRJNA504891).

We used the first three datasets to evaluate single-sample and multi-sample binning mode and the last two human gut projects as hold-out datasets to evaluate the pretrained model in SemiBin.

For simulated datasets, we used the gold standard contigs provided as part of the challenge. For real datasets (except PRJNA504891), short reads were first filtered with NGLess (Coelho et al., 2019) (version 1.0.1, default parameters) to remove low-quality reads and host-matching reads (human reads for human gut datasets, dog reads for dog gut datasets). These pre-processed reads were then assembled using Megahit (Li et al., 2015) (version 1.2.4, default parameters) to assemble reads to contigs. For PRJNA504891, we used Megahit (Li et al., 2015) (version 1.2.8, default parameters) to assemble reads to contigs. We mapped reads to the contigs with Bowtie2 (Langmead and Salzberg, 2012) (version 2.4.1, default parameters) to generate the BAM files used in the binning. For multi-sample binning mode, contigs from all samples were concatenated together. Then reads from every sample were mapped to the concatenated contig to get the BAM files for every sample.

### 2.3 Methods included in the benchmarking

We compared SemiBin to other methods in three binning modes. For single-sample and co-assembly binning of CAMI I datasets, we compared our method to the following methods: Maxbin2 (version 2.2.6) (Wu et al., 2016), Metabat2 (version 2) (Kang et al., 2019), VAMB (latest version) (Nissen et al.), COCACOLA (latest version) (Lu et al., 2017), SolidBin (version 1.3) (Wang et al., 2019). SolidBin is the only existing semi-supervised binning method and it has different modes. Here, we focused on the comparison of modes with information from reference genomes: SolidBin-coalign (which generates *must-link* constraints from reference genomes), SolidBin-CL (which generates *cannot-link* constraints from reference genomes) and SolidBin-SFS-CL (which generates *must-link* constraints from feature similarity and reference genomes). We also added SolidBin-naive (without additional information) to show the influence of different semi-supervised modes.

For multi-sample binning of CAMI II datasets, we compared to the existing multi-sample binning tool VAMB, which clusters concatenated contig based on co-abundance across samples and then split the clusters according to original samples and default Metabat2. For more comprehensive benchmarking, we converted Metabat2 to multi-sample mode. We used *jgi_summarize_bam_contig_depths* (with default parameters) to calculate depth values using the BAM files from every sample mapped to the concatenated contig. Then, we ran Metabat2 to bin contigs for every sample with abundance information across samples, which is the similar idea of the multi-sample mode in SemiBin.

We also benchmarked single-sample and multi-sample binning mode in real datasets. For single-sample binning, we compared the performance of SemiBin to Maxbin2, Metabat2 and VAMB, and for multi-sample binning, we compared to VAMB. These tools have been shown that they can perform well in real metagenomic projects (see Supplementary Table 3). For VAMB, we set the minimum length as 2,000bp. For SolidBin, we ran SolidBin with constraints generated from annotation results with MMseqs2 (Steinegger and Söding, 2017; Mirdita et al., 2020) and we used the binning results after the postprocessing with CheckM. For SemiBin, we ran the whole pipeline described in the Methods (with default parameters, except for the inclusion of the reclustering step). For other benchmarking methods, we ran tools with default parameters.

To evaluate the effectiveness of the semi-supervised learning in SemiBin, we also benchmarked modified versions of SemiBin:

1. NoSemi, where we removed the semi-supervised learning step in the SemiBin pipeline, and cluster based on the original inputs.
2. SemiBin_m, where we removed the semi-supervised learning step, but directly used the *must-link* constraints in the sparse graph (by adding a link between *must-link* contigs).
3. SemiBin_c was analogous to SemiBin_m, but we used *cannot-link* constraints by removing links in the sparse graph between *cannot-link* contigs.
4. SemiBin_mc combines the operations of SemiBin_m and SemiBin_c. Then, extracting bins from the constructed graph as in the standard pipeline was performed.

### 2.4 Timing evaluations

For evaluating the computational resource usage, we used two Amazon Web Services (AWS) virtual machines. We used the machine type *ga4d.x4large* to run SemiBin in CPU-mode. This machine contains 2nd generation AMD EPYC processors, 16 cores and 64 GB RAM memory. Additionally we used the type *g4dn.4xlarge* to run SemiBin in GPU-mode. This machine contains an NVIDIA Tesla T4 GPU.

### 2.5 Evaluation metrics

For simulated dataset in CAMI, we used AMBER (version 2.0.1) (Meyer et al., 2018) to calculate the completeness (recall), purity (precision), and F1-score to evaluate the performance of different methods.

In the real datasets, as ground truth is not available, we evaluated the completeness and contamination of the predicted bins with CheckM (Parks et al., 2015) (version 1.1.3, using *lineage_wf* workflow with default parameters). We defined high-quality bins as those with completeness > 90%, contamination < 5% (Bowers et al., 2017) and also passing the chimeric detection implemented in GUNC (Orakov et al., 2021) (version 0.1.1, with default parameters). Medium-quality bins are defined as completeness ≥ 50% and contamination < 10% and low-quality bins are defined as completeness < 50% and contamination < 10% (Bowers et al., 2017).

### 2.6 Model transfer between environments

To evaluate the generalization of the learned models, we selected three models as training sets from human gut, dog gut, and ocean microbiome datasets. In each dataset, we selected a model from the sample that could generate the highest number, median number, and lowest number of high-quality bins. For the human gut dataset, we termed them as human_high, human_median and human_low, with models from the other environments named analogously. For every environment, we also randomly sampled 10 samples from the rest of the samples as testing sets (no overlap in the training sets and testing sets). Then, we transferred these models to the testing sets from the same environment or different environments and used the embeddings from these models to bin the contigs.

To evaluate the effect of training with different numbers of samples, for every environment, we also randomly chose 10 samples as testing sets and trained the model on different number of training samples (randomly chosen 1, 3, 5, 10, 15, 20 samples; no overlap in training sets and testing sets). For each number of samples, we randomly chose samples and trained the model 5 times.

To evaluate the pretraining approach, we used another two human gut datasets. We termed SemiBin with a pretrained model from the human gut dataset used before as SemiBin(pretrain; external). We also trained a model from 20 randomly chosen samples from the hold-out datasets and applied it to the the same dataset; an approach we termed SemiBin(pretrain; internal). We benchmarked SemiBin(pretrain; external), Metabat2, original SemiBin, and SemiBin(pretrain; internal).

### 2.7 Metagenome-assembled genomes (MAG) analyses

To identify the overlap between the bins from SemiBin and Metabat2 with single-sample binning, we used Mash (version 2.2, with default parameters) (Ondov et al., 2016) to calculate the distance between bins from two methods. Then, we assigned corresponding bins with Mash distance ≤0.01 and we considered these two bins as the same genome in subsequent comparisons. After obtaining the overlap of bins sets, we classified the high-quality bins from one method into 4 classes: HQ-HQ: also high-quality in the other method; HQ-MQ: medium-quality in the other; HQ-LQ: low-quality or worse in the other; and HQ-Miss: could not be found in the other. We calculated the recall, precision, and F1-score for the HQ-HQ component. Recall and precision is completeness and 1 - contamination estimated from CheckM. F1-score is 2 ×(recall × precision)/(recall + precision). To evaluate the species diversity of different methods, we annotated high-quality bins with GTDB-Tk(version 1.4.1, using *classify_wf* workflow with default parameters) (Chaumeil et al., 2020).

For the analysis of *Bacteroides vulgatus* bins, average nucleotide identity (ANI) comparisons were calculated using fastANI (Jain et al., 2018) (version 1.32, *–fragLen 1000*). *B. vulgatus* bins were annotated with Prokka(version 1.14.5, with default parameters) (Seemann, 2014). Pan-genome analyses were carried out using Roary(version 3.13.0, *-i 95 −cd 100*) (Page et al., 2015). We used Scoary(version 1.6.16, *-c BH*) (Brynildsrud et al., 2016) to identify genes with significant differences in the human and dog gut microbiome datasets. Phylogeny reconstructions of core genes were performed with IQTREE using 1000 bootstrap pseudoreplicates for model selection (version 1.6.9, *-m MFP −bb 1000 -alrt 1000*) (Nguyen et al., 2015), and visualized with ggtree package(v1.8.154) (Yu et al., 2017). Principal component analysis (PCA) was done using the *prcomp* function from the *stats* (Team et al., 2013) package.

## 3 Data availability

The sequence data used in the study are publicly available with study accessions PRJEB27928, PRJEB20308, PRJEB1787, PRJNA504891 and PRJNA290729. The simulated CAMI I(low, medium and high complexity) and CAMI II datasets(skin and oral cavity from Toy Human Microbiome Project Dataset) can be downloaded from https://data.cami-challenge.org/participate. The MAGs that generated from real metagenomes in the benchmarking can be obtained from Human gut MAGs, Dog gut MAGs and Ocean MAGs.

## 4 Code availability

Code for the tool can be found on GitHub at https://github.com/BigDataBiology/SemiBin/ and is freely available under the MIT license. The analysis code and intermediate results can be found on Github at https://github.com/BigDataBiology/SemiBin_benchmark.

## 5 Acknowledgements

This research was supported by the National Natural Science Foundation of China (grant 31950410544, LPC), the Shanghai Municipal Science and Technology Major Project (grant 2018SHZDZX01, XMZ and LPC) and ZHANGJIANG LAB (XMZ and LPC).

We thank Anthony Fullam and Thomas Sebastian B. Schmidt (Bork group, European Molecular Biology Laboratory) for providing us with the assembled contigs for the metagenomes used (with the exception of those from PRJNA504891). We thank members of the Coelho and Zhao groups for their comments and discussions throughout the development of the project. We also thank Marija Dmitrijeva (University of Zurich) for helpful comments on a previous version of the manuscript. Beta users of SemiBin are thanked for their suggestions and bug reports.

## 6 Author information

### 6.2 Contributions

L.P.C conceived the study and supervised the project. S.P, X.Z and L.P.C designed the method. S.P. and L.P.C wrote the software. S.P, C.Z and L.P.C designed and performed the analyses. S.P. wrote the first draft of the manuscript. All authors contributed to the revision of the manuscript prior to submission and all authors read and approved the final version.

## 7 Supplementary Text

### 7.1 Noise and bias of constraints generation from taxonomic contig annotations

Following taxonomic annotation of contigs, *cannot-link* constraints between contigs were automatically extracted (see Methods). We also attempted to extract *must-link* annotations by defining pairs of contigs with same annotations at species level (with scores both above 0.95) as possessing a *must-link* constraint between them. In the CAMI simulated data, it is possible to evaluate the accuracy and coverage of these annotations. We observed that *cannot-link* constraints had a very high accuracy and, thus, they are used for training (see Supplementary Table 2). For *must-link* constraints, there are two issues. First, for most situations, the accuracy of *must-link* constraints was low, which would lead to noise in the training. Second, the *must-link* constraints only covered a small part of the genomes in the environment and it would lead to bias of the learning of model. Owing to the noise and bias of the *must-link* constraints obtained from taxonomic annotations, we chose to not use them and instead to generate *must-link* constraints by breaking up long contigs.

### 7.2 Evaluation of the learning ability of SemiBin

To evaluate the learning ability of SemiBin, namely that it has the ability to learn the underlying genome structure from the *must-link* and *cannot-link* constraints, not just reproduce its inputs, we compared the full pipeline to NoSemi (no constraints are used) as well as SemiBin_m, SemiBin_c and SemiBin_mc which directly use the *must-link* and *cannot-link* constraints to generate the graph without semi-supervised learning step (see Methods). The complete SemiBin pipeline performed similarly or better (average 12.4% more high-quality bins) in low and medium complexity datasets and large improvements (average 34.6% more high-quality bins) in high complexity dataset (see Supplementary Fig. 11) compared to the second best binner. SemiBin could also reconstruct on average 7.0% and 16.0% more distinct high quality strains in Skin and Oral datasets, respectively (see Supplementary Fig. 12). These results show the ability of semi-supervised model in learning the genome structure of the environment beyond what was present in the inputs, especially in complex environments.

### 7.3 Robustness of results to changes in the *max-edges* parameter

We compared the parameter *max_edges* in Metabat2 and SemiBin on CAMI I datasets. For both tools, this *max_edges* parameter controls the number of edges of each node (each contig) in the graph that will be considered in binning. Both SemiBin and Metabat2 were robust to the setting of *max_edges*. Nonetheless, SemiBin could reconstruct average 26.0%, 11.9% and 36.4% more high-quality bins (see Supplementary Fig. 6) than Metabat2.

Compared to NoSemi with different *max_edges* parameters, SemiBin was more robust as the results of NoSemi deteriorated with increasing number of edges (see Supplementary Fig. 10). This also indicates the better embedding obtained from the deep learning.

### 7.4 Evaluation of the performance of SemiBin in simulated datasets

We trained the deep learning model on the CAMI low-complexity dataset and compared to semi-supervised SolidBin-coalign and SolidBin-CL. The original features and embedded ones from SolidBin-coalign, SolidBin-CL and SemiBin were visualized for every genome using t-SNE (Maaten and Hinton, 2008). Semi-supervised deep learning in SemiBin led to a better separation between genomes and better aggregation within genomes (see Fig. 2d and Supplementary Fig. 7).

When comparing the different versions of SolidBin on CAMI I datasets, in most situations, SolidBin with additional information performed worse than SolidBin-naive which showed that the semi-supervised Ncut algorithm used in SolidBin could not leverage additional information very well, perhaps due to the noise in these annotations (see Supplementary Table 2).

### 7.5 Applying SemiBin to real data

In the human gut dataset, SemiBin with multi-sample binning reconstructed more high-quality bins than SemiBin with single-sample binning, but these came from fewer distinct species, genera and families. This showed that multi-sample binning (which uses abundance across several samples) might lead to the recovery of more genomes from species occurring in multiple samples, while overlooking rare species (see Supplementary Fig. 16).

By default, SemiBin learns an embedding model for each sample. To evaluate the generalization of this learned model, we transferred models between different samples and environments. For this, we selected three models from each environment studied (termed as high, median and low according to the number of high-quality bins they reconstructed, see Methods) and tested their performance on 10 randomly selected samples from each environment, using single-sample binning (no overlap in training data and testing data, see Methods). Not unexpectedly, training a new model for every sample resulted in the highest number of high-quality bins, while SemiBin with a pretrained model from the same environment achieved the second best result (see Supplementary Fig. 13). Nonetheless, it is noteworthy that SemiBin with a pretrained model from the same environment could still perform better than Metabat2, reconstructing at most 26.0%, 59.2% and 60.0% more high-quality bins on human gut, dog gut, and ocean testing datasets, respectively. In most situations, the pretrained model that generated the highest number of high-quality bins performed better than the model that generated median and lowest number from the same environment. Furthermore, transferring models between different environments could still improve results compared to NoSemi version of SemiBin and in some situations, transferring across environments performed better than Metabat2 (see Supplementary Fig. 13).

The results of model transfer indicated that the semi-supervised model learned high-level or shared structure of microorganism between environments. However, transferred models still underperformed models learned for each sample and there was a dependency on the sample used to train. Thus, we attempted to mitigate this by learning models on multiple samples simultaneously. This approach achieved the best results, while not requiring computationally-costly per-sample training (see Fig. 2 and main text).

### 7.6 Comparison of contig taxonomic annotation methods

We compared the results of SemiBin with two different contig taxonomic annotation tools: CAT (von Meijenfeldt et al., 2019) and MMseqs2 (Steinegger and Söding, 2017; Mirdita et al., 2020) on CAMI I simulated datasets and human gut, dog gut microbiome real datasets, evaluating the methods by the final number of high-quality bins recovered (see Supplementary Fig. 20). Besides internal algorithmic differences, CAT annotates contigs to the NCBI taxonomy, while with MMseqs2 we used the GTDB as the target. Interestingly, in simulated datasets, CAT performed slightly better than MMseqs2; while in real datasets, MMseqs2 performed significantly better. This illustrates the perils of over-reliance on simulated benchmarks, which do not capture all the complexity of real data. Therefore, we chose MMseqs2 and the GTDB as the default contig annotation tool and taxonomy in SemiBin (although it can easily be replaced by the user).

### 7.7 Analysis of *Bacteroides vulgatus* strains

Comparing the average nucleotide identity (ANI) of 50 *B. vulgatus* bins (49 recovered from our data, and one reference genome GCF000012825), we found that the ANI of *B. vulgatus* within dog gut microbiome (average 99.4%) was significantly higher than those within human gut microbiome (average 98.8%) (see Supplementary Fig. 21a). To further explore the diversity of the *B. vulgatus*, we predicted and annotated protein-coding sequences (CDS) with Prokka (Seemann, 2014) and built the pangenome with Roary (Page et al., 2015) using a threshold of 95% (nucleotide identity) for gene similarity. We identified a total of 15,382 genes, of which only 458 are core genes shared between all 50 MAGs. These core genes were used to infer a maximum likelihood phylogenetic tree using IQTREE (Nguyen et al., 2015) (see Fig. 2d). It was clear that MAGs from the dog gut tend to cluster together. The same results can be found when performing principal component analysis (PCA) based on the presence/absence of genes of the 50 *B. vulgatus* strains (see Supplementary Fig. 21b).

We observed that a total of 13 genes were differentially present in dog and human gut strains (see Fig. 2 and main text). Using the Global Microbial Gene Catalog (GMGC, https://gmgc.embl.de/) for external validation (GMGC contains 762 *B. vulgatus* high-quality strains from human gut, recovered using Metabat2), we verified that 7 of them were still significant (see Supplementary Fig. 22). Furthermore, in the maximum likelihood phylogenetic tree including GMGC strains, MAGs from dog guts still clustered together (see Supplementary Fig. 22).

---

**Supplementary Fig 1.**
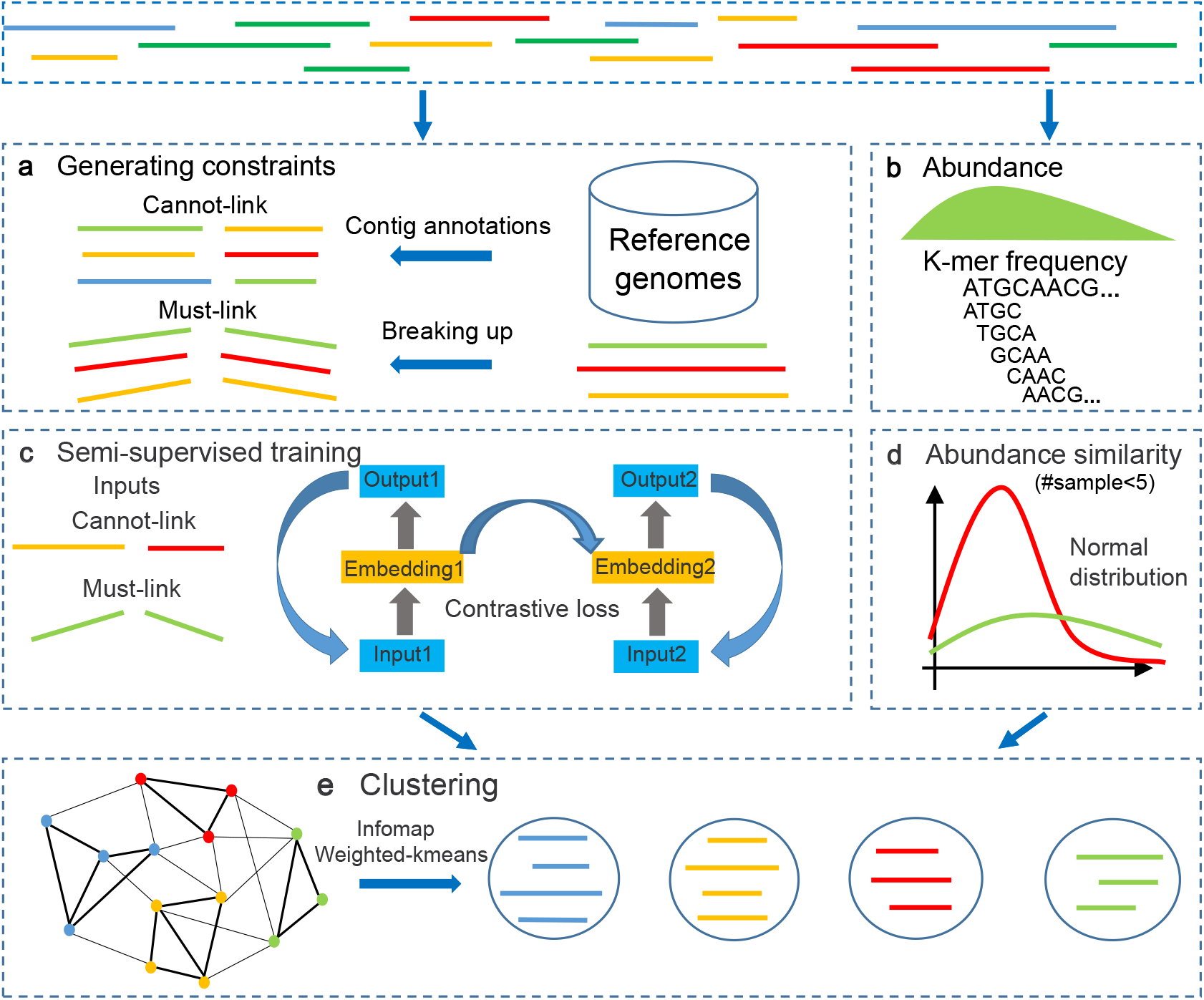
Overview of the SemiBin pipeline. **a,**Generating *must-link* constraints by breaking up contigs artificially and generating *cannot-link* constraints based on contig taxonomic annotations (*i.e.* GTDB reference genomes). **b,** Calculating abundance value (average and variance of the number of reads per base) and *k*-mer frequency of every contig. **c,** Training semi-supervised siamese neural network of the *cannot-link* and *must-link* constraints as inputs. The learned embedding will be used in step **e** for binning. textbfd, Based on the assumption that the number of reads per base obeys normal distribution, calculating the Kullback-Leibler divergence of the normal distributions of two contigs. SemiBin uses this value as the abundance similarity when the number of samples used is smaller than 5. **e,** Generating a sparse graph with the embedding distance and abundance similarity and Infomap algorithm is used to get the preliminary bins. Finally, SemiBin uses weight *k*-means to recluster bins whose mean number of single copy genes is greater than one to get the final bins.

**Supplementary Fig 2.**
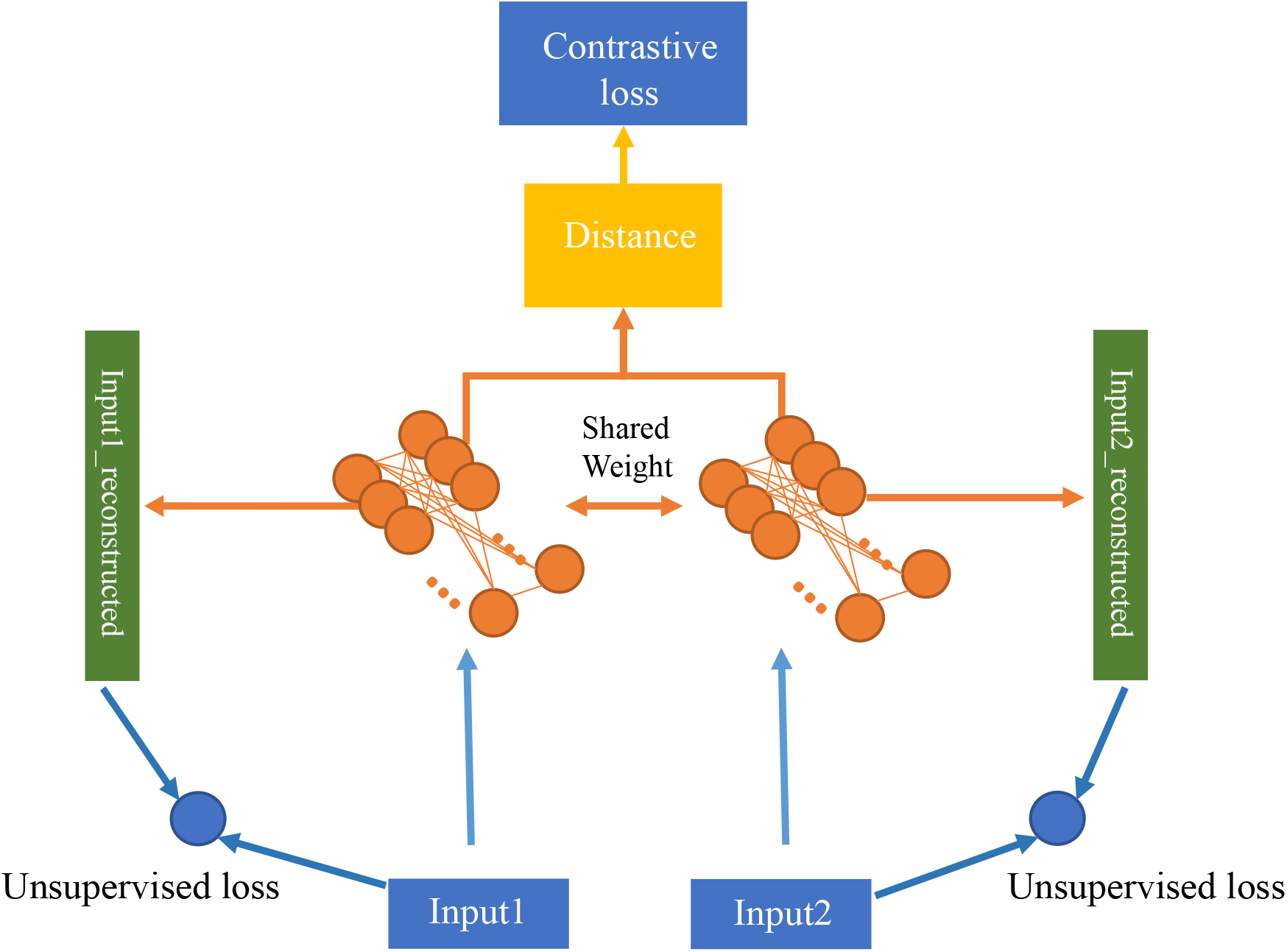
Semi-supervised neural network model used in the SemiBin. A pair (input1, input2) is input to a shared-weight neural network. Input1 can be *k*-mer frequencies and the abundance distribution (n ≥ 5) or just the *k*-mer frequencies (n < 5). During the training, the unsupervised loss and the contrastive loss are optimized at the same time. After the training, the embedding of the inputs can be used for the following binning.

**Supplementary Fig 3.**
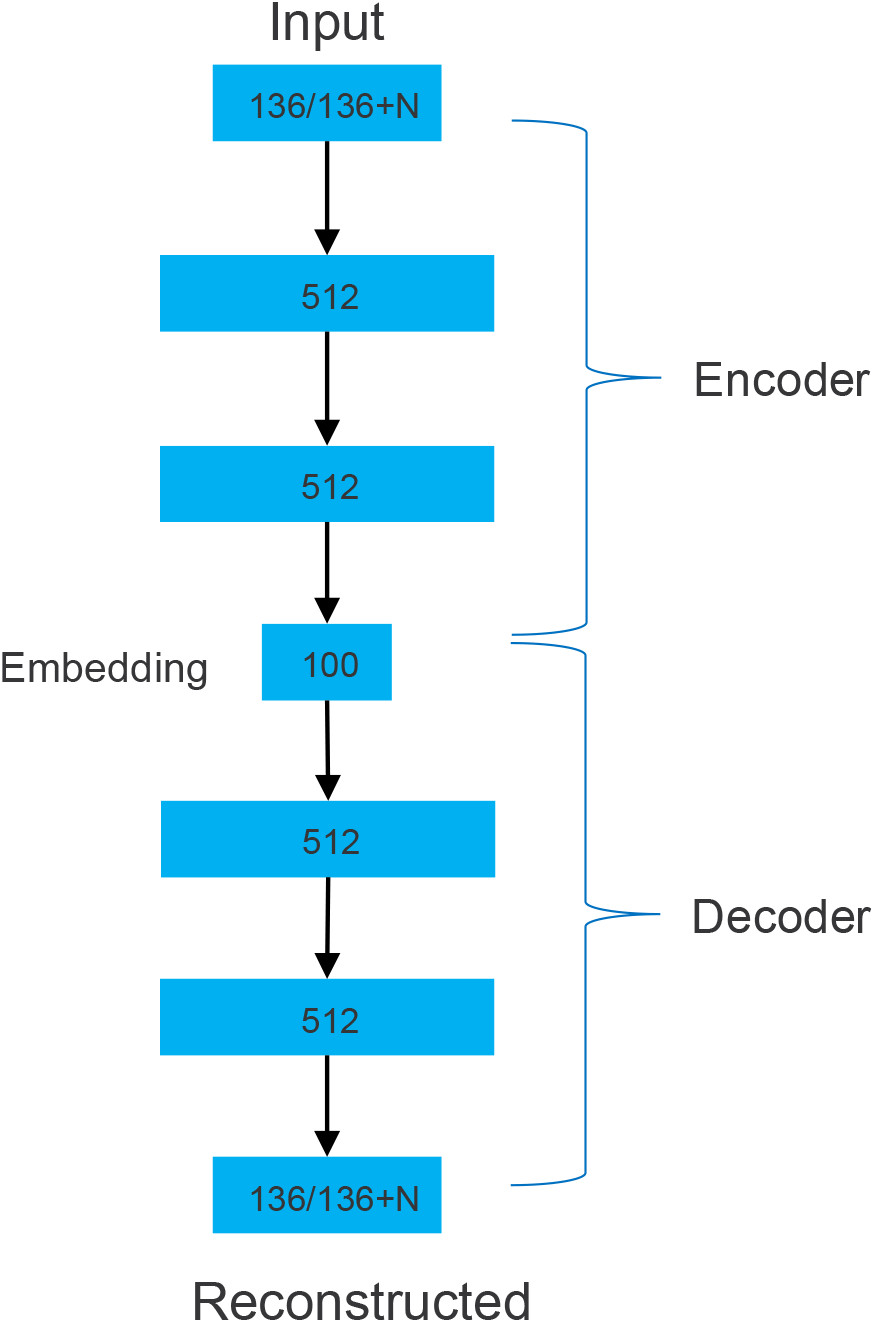
Structure of the semi-supervised neural network model used in the SemiBin. The number in the box is the number of neurons in every layer. The neural network used in the SemiBin is a shared-weight autoencoder. An encoder network encodes the inputs to 100 dimension features and a decoder network reconstructs the original inputs. After the training, the 100 dimensions embeddings are used in the binning.

**Supplementary Fig 4.**
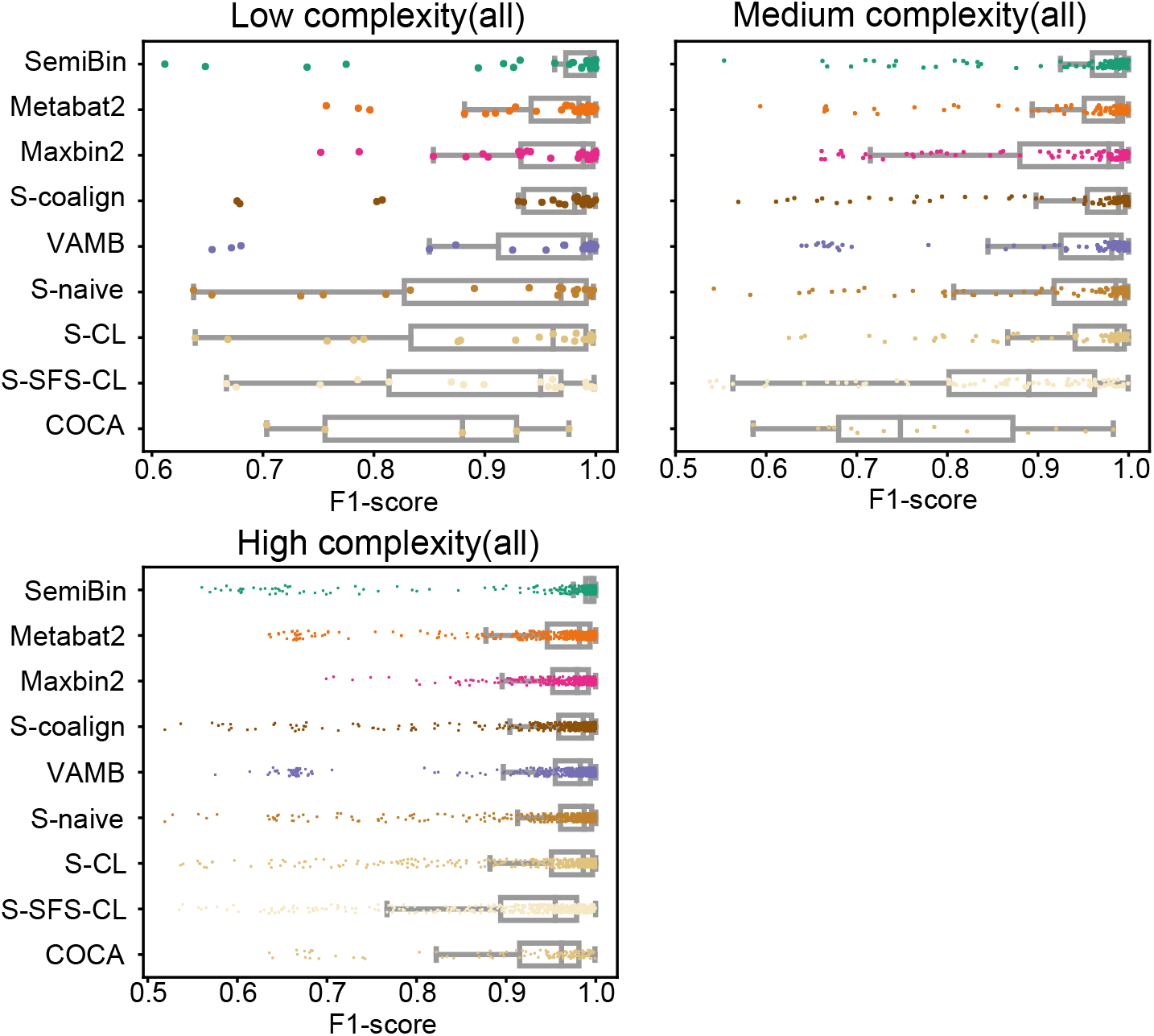
SemiBin outperformed other binners on CAMI I datasets with single-sample and co-assembly binning. Shown is F1-score distribution of bins (completeness ≥ 50%; contamination ≥ 50%) reconstructed from low, medium and high complexity datasets of CAMI I with single-sample and co-assembly binning. (Top to down are the results of SemiBin, Metabat2, Maxbin2, SolidBin-coalign, VAMB, SolidBin-naive, SolidBin-CL, SolidBin-SFS-CL and COCACOLA).

**Supplementary Fig 5.**
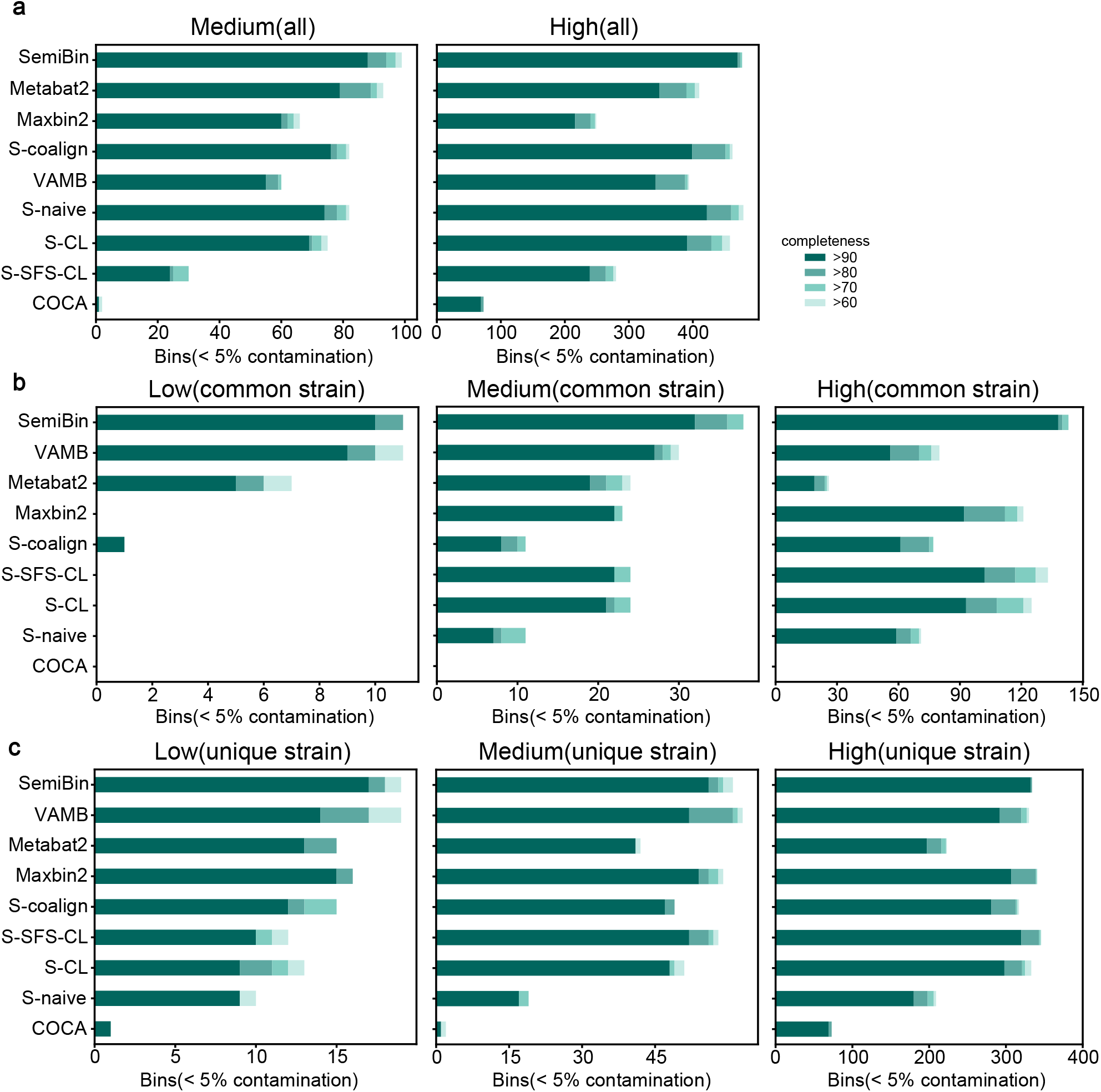
SemiBin outperformed other binners in CAMI I datasets. Genomes in CAMI I datasets are defined as common strains and unique strains according to the genome similarity. Common strains are defined as genomes with an ANI (average nucleotide identity) ≥ 95% to the most similar genomes in the environment and unique strains are defined as genomes with < 95% ANI value to any other genome. Shown is the number of reconstructed genomes per method above certain completeness and contamination < 5% for **a,** medium and high complexity datasets considering all strains; **b,** three datasets considering common strains; and **c,** three datasets considering unique strains. SemiBin reconstructed more high-quality bins (considering all strains, common strains and unique strains), especially for common strains which is a big challenge for binning in an environment with multiple strains.

**Supplementary Fig 6.**
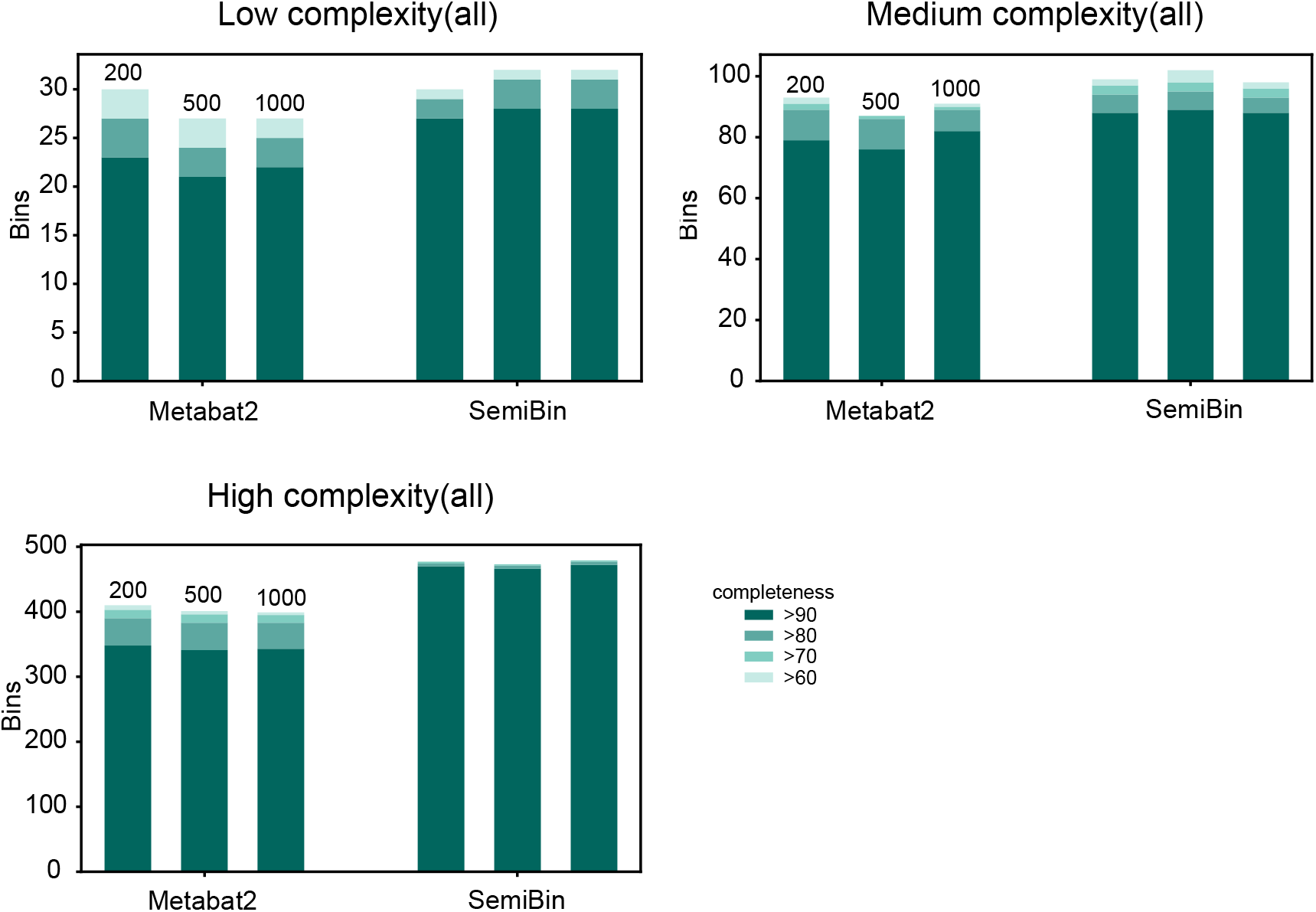
SemiBin outperformed Metabat2 with different *max_edges* settings (200, 500 and 1000) in CAMI I datasets. We investigated the influence of different values (200, 500, 1000) of the same parameter *max_edges* used in SemiBin and Metabat2. SemiBin outperformed Metabat2 with all situationns. **a,** low complexity dataset; **b,** medium complexity dataset; and **c,** high complexity dataset.

**Supplementary Fig 7.**
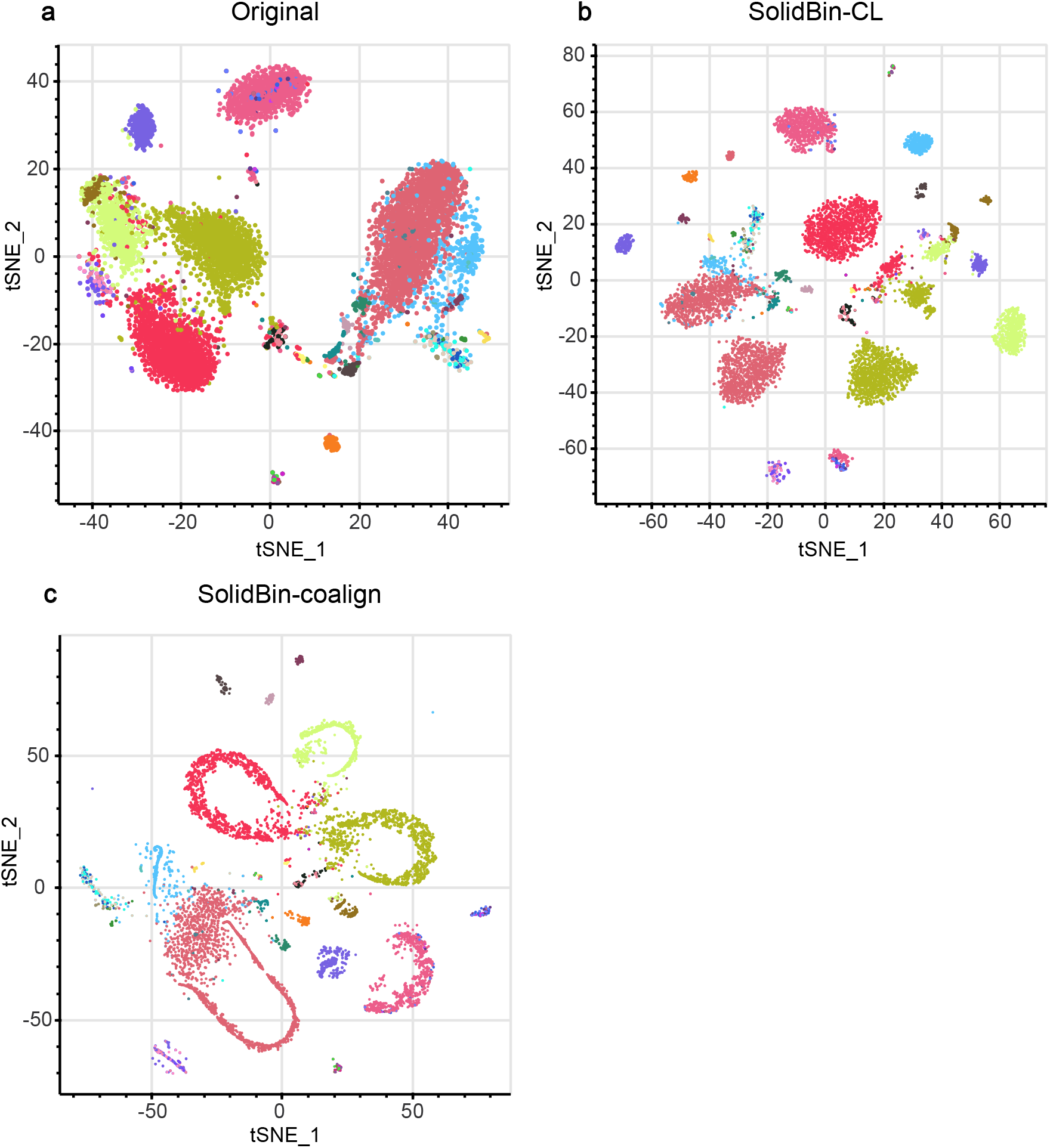
Visualization of original features and embeddings from SolidBin in CAMI I low complexity dataset. Shown are **a,** the original features and embeddings from **b,** SolidBin-CL and **c,** SolidBin-coalign(two versions of SolidBin that used additional information from reference genomes) with t-SNE on the 40 genomes from CAMI I low complexity dataset.

**Supplementary Fig 8.**
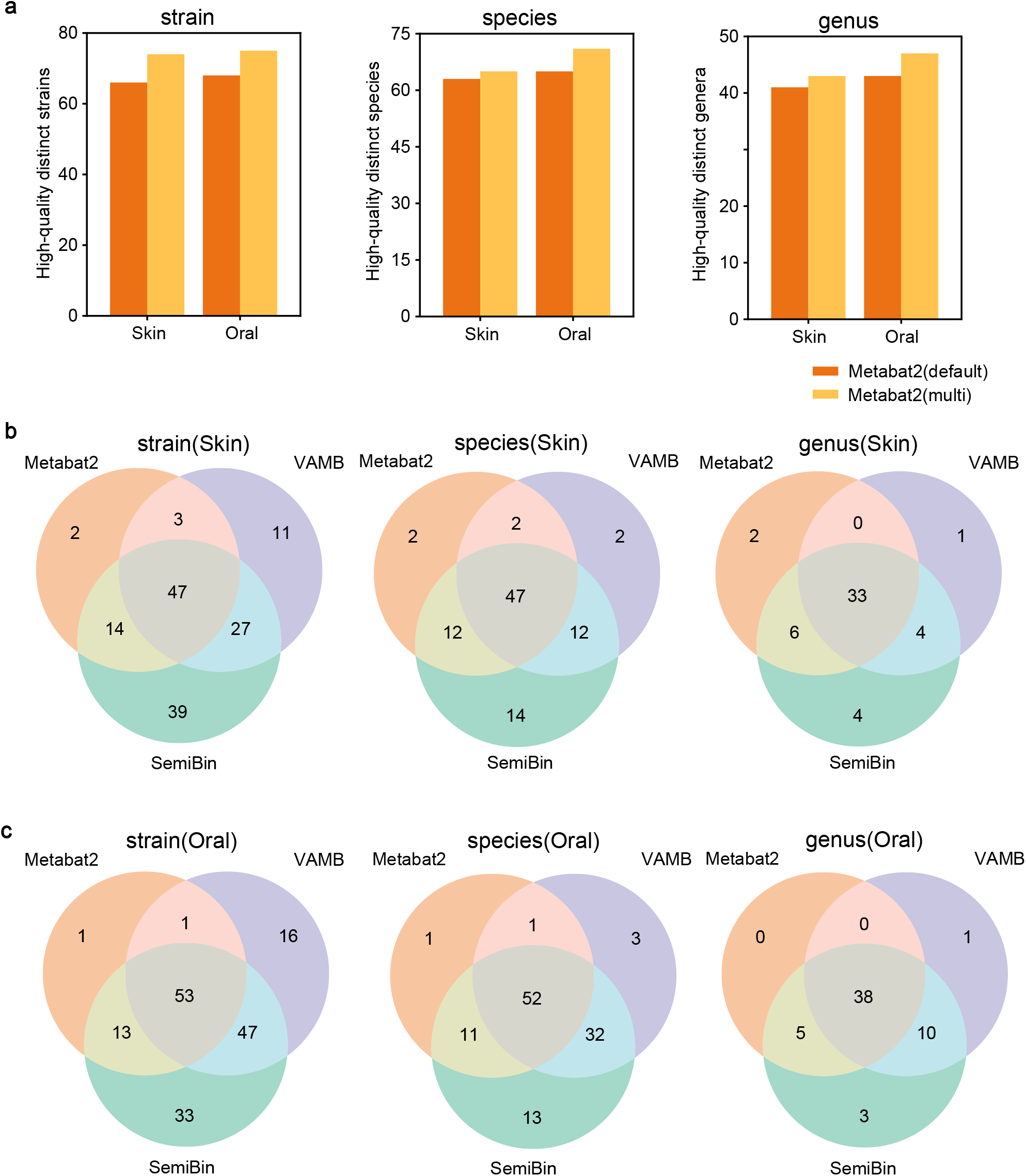
SemiBin outperformed Metabat2 and VAMB in CAMI II datasets with multi-sample binning. **a,**The comparison of Metabat2 with single-sample binning (Metabat2(default)) and our adapted multi-sample binning (Metabat2(multi)). The adapted multi-sample binning Metabat2 led only modest improvements. **b,** and **c,** Shown are the overlaps of the reconstructed distinct high quality strains, species and genus for the Skin and Oral datasets. SemiBin reconstructed more distinct strains, species and genera compared to VAMB and Metabat2.

**Supplementary Fig 9.**
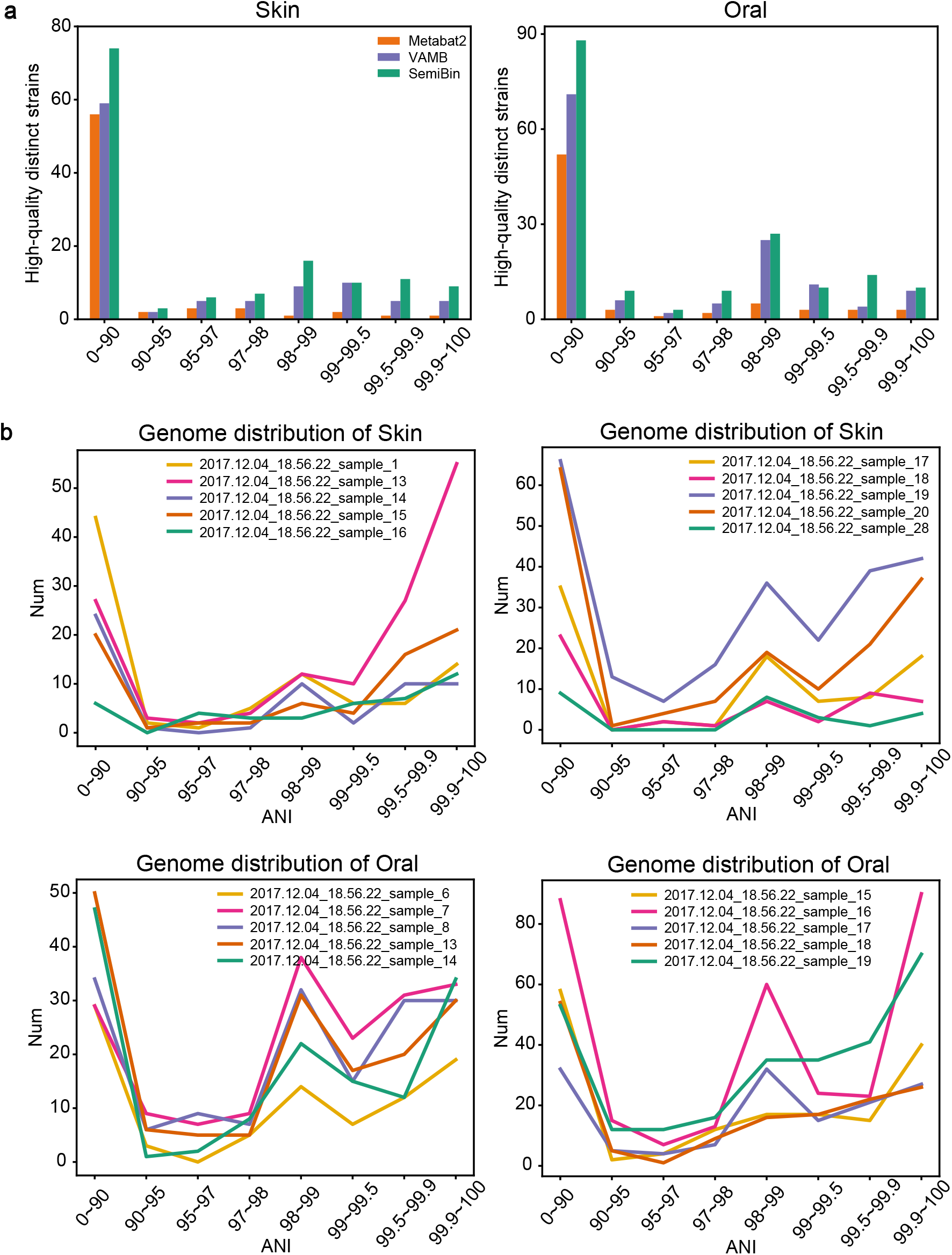
SemiBin outperformed Metabat2 and VAMB across almost all ANI intervals in CAMI II datasets. We stratified the datasets according to the ANI value which is defined as the ANI of one genome to the most similar genome in the same sample. We calculated the number of distinct high-quality strains in every interval. Shown are *a,* the number of reconstructed distinct high-quality strains in every interval, and *b,* the genome distribution according to the ANI values for every sample of Skin and Oral datasets.

**Supplementary Fig 10.**
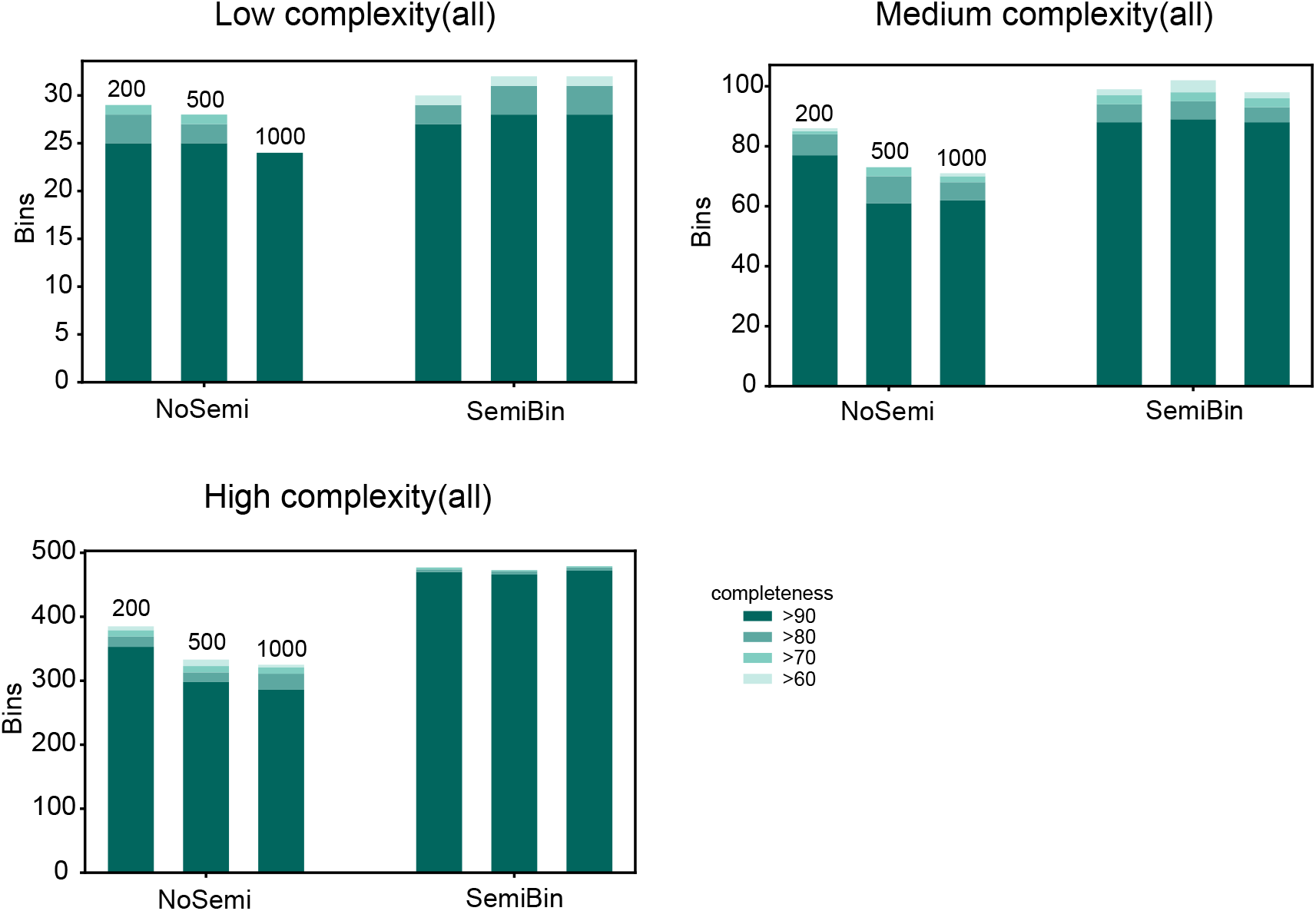
Semi-supervised deep learning in SemiBin significantly improved binning results. We compared SemiBin to the NoSemi version (removing the semi-supervised learning part in SemiBin) to show the performance of the semi-supervised learning used in SemiBin. Shown is the number of reconstructed genomes (with certain completeness and smaller than 5% contamination) for **a,** low complexity dataset; **b,** medium complexity dataset; and **c,** high complexity dataset.

**Supplementary Fig 11.**
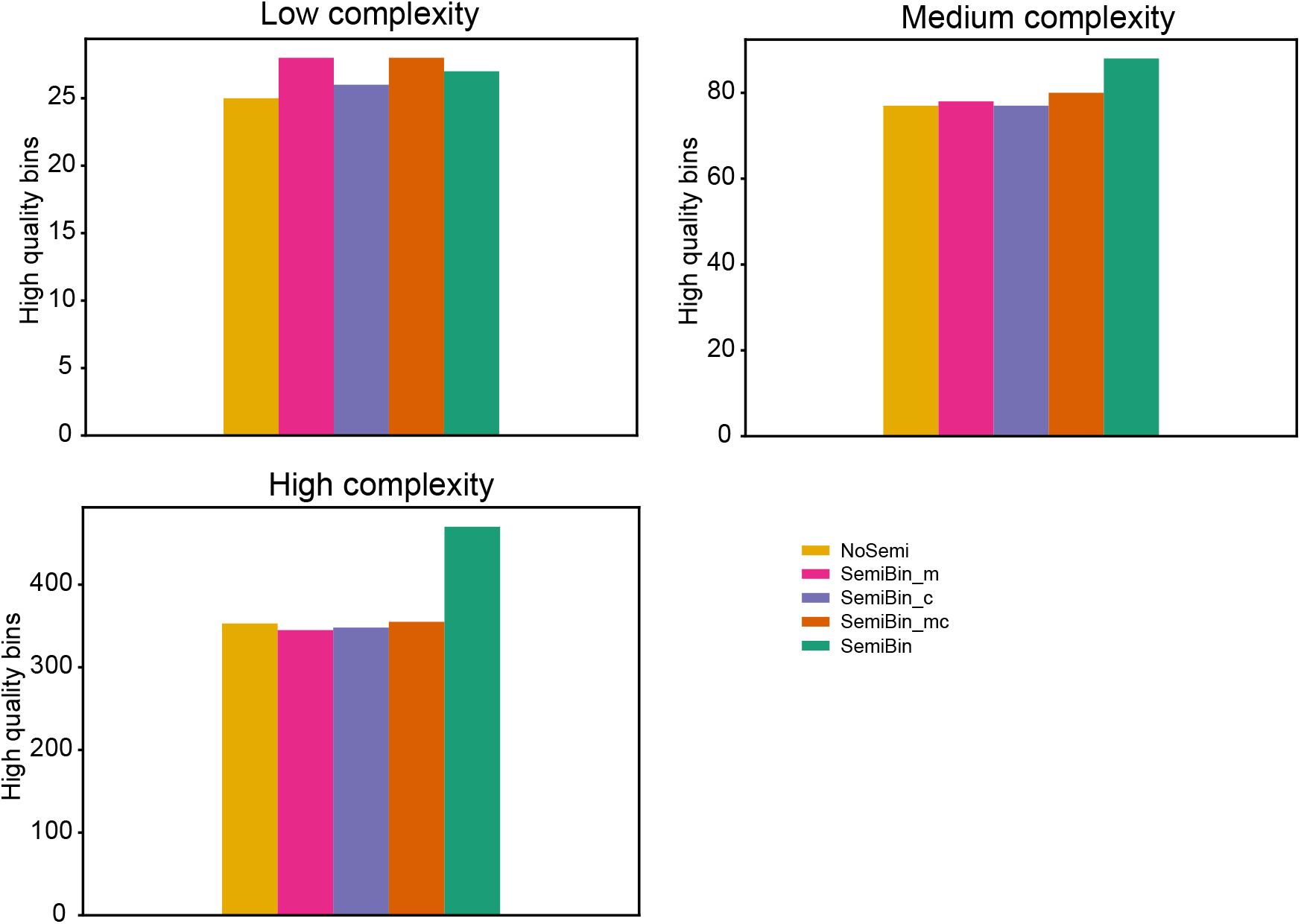
Semi-supervised deep learning in SemiBin learnt the underlying structure of the environment in CAMI I datasets. To evaluate that the semi-supervised learning model could learn the underlying structure of the environment (not just remembering the *must-link* and *cannot-link* constraints), we compared SemiBin to the NoSemi version (removing the semi-supervised learning part in SemiBin), SemiBin_m (directly using *must-link* constraints to generate the sparse network for clustering, no semi-supervised learning), SemiBin_c (directly using *cannot-link* constraints to generate the sparse network for clustering, no semi-supervised learning), and SemiBin_mc (directly using *must-link* and *cannot-link* constraints to generate the sparse network for clustering, no semi-supervised learning) (see Methods). Shown is the number of high-quality bins in low complexity, medium complexity and high quality datasets from CAMI I.

**Supplementary Fig 12.**
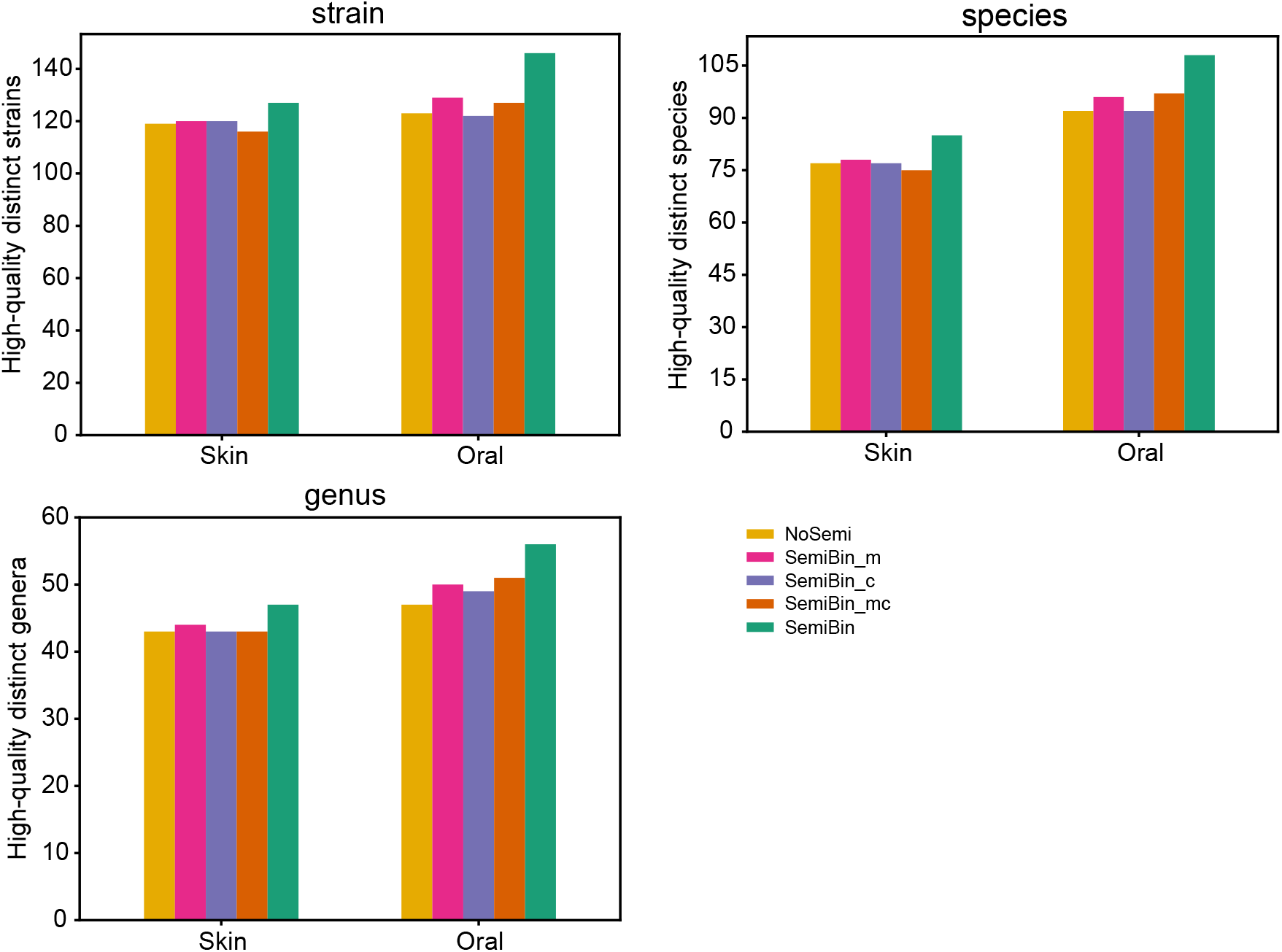
Semi-supervised deep learning in SemiBin learnt the underlying structure of the environment in CAMI II datasets. Shown is the number of high-quality distinct strains, species and genera in Skin and Oral datasets. The methods used here are the same as those in Supplementary Fig. 11.

**Supplementary Fig 13.**
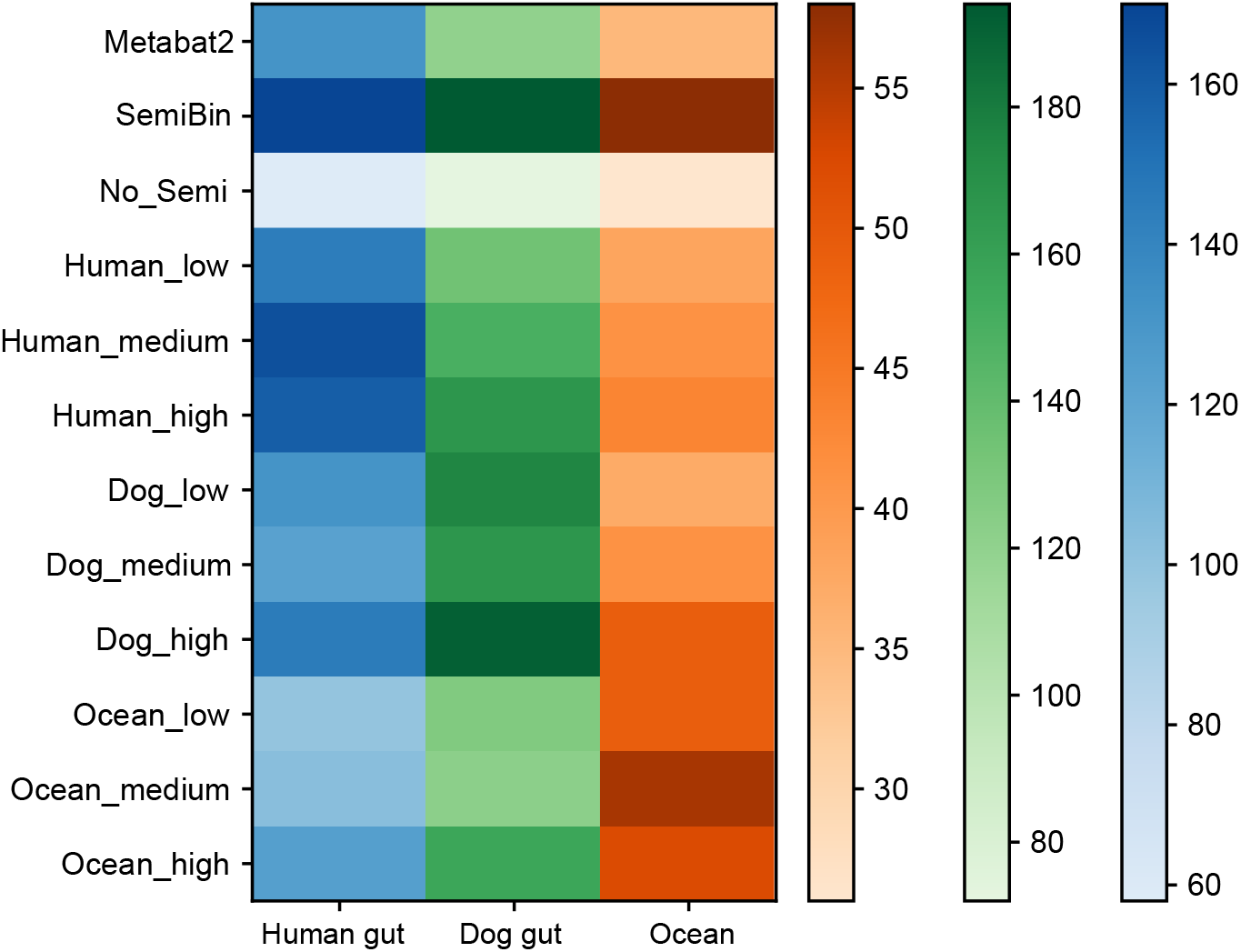
SemiBin with pretrained model trained from one sample got competitive binning results. We transferred the learned semi-supervised models from one sample between human gut, dog gut and ocean datasets. We chose three models from samples which reconstructed the highest, median and lowest number of high-quality bins for each environment, and termed these models as human_high, human_medium, human_low, dog_high, dog_medium, dog_low, ocean_high, ocean_medium, ocean_low. For every environment, we randomly chose 10 samples which were not used before as testing sets (No overlap in training samples and testing samples). The transferring results compared to Metabat2, original SemiBin and nosemi version of SemiBin(NoSemi) are shown as the number of high-quality bins for each environment (the darker the color, the higher number of high-quality bins).

**Supplementary Fig 14.**
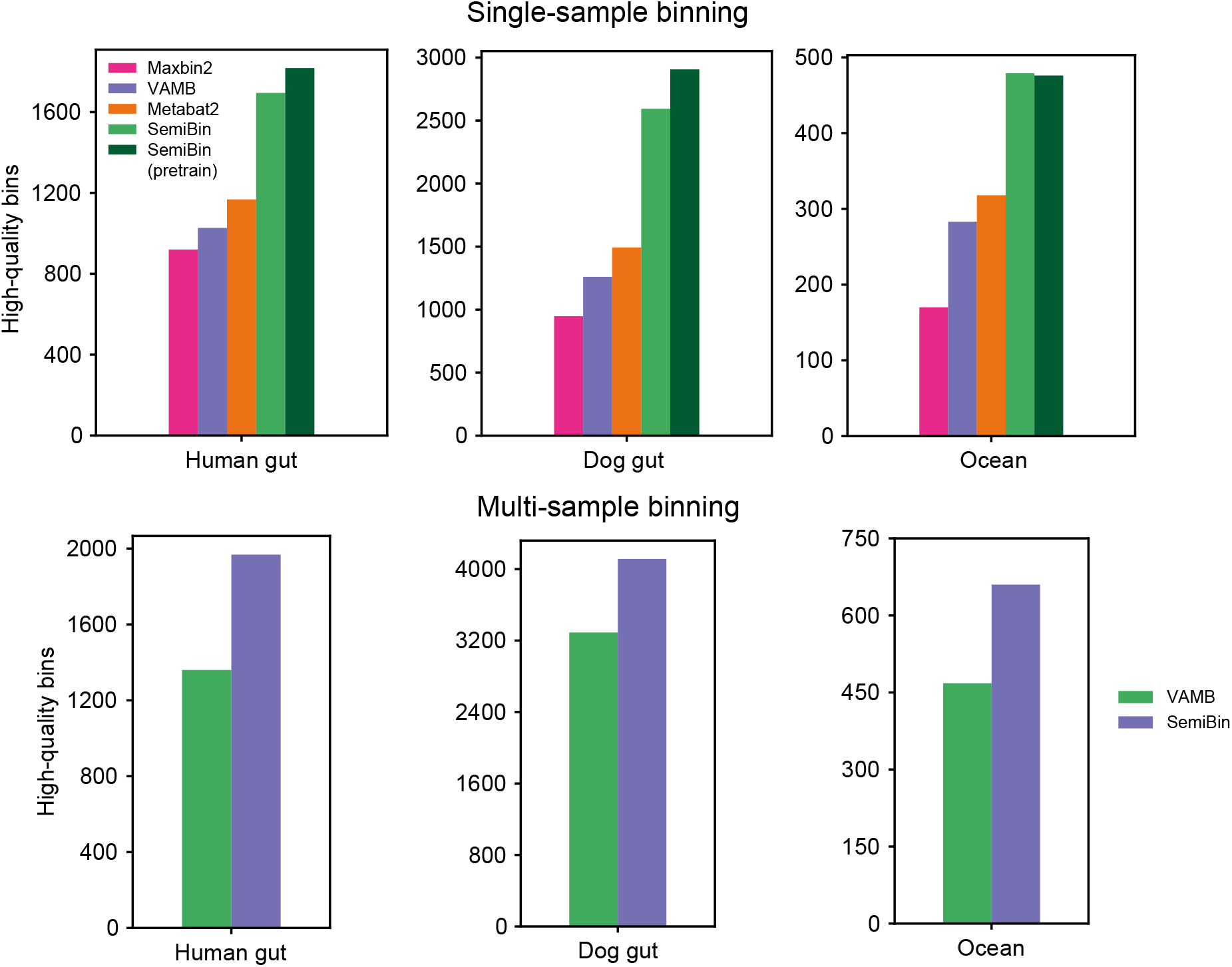
SemiBin outperformed other binners in real datasets with single-sample and multi-sample binning evaluated by CheckM. The high-quality bin is defined as a bin with completeness > 90% and contamination < 5% evaluated by CheckM. Shown is the number of high-quality bins generated by Maxbin2, VAMB, Metabat2, SemiBin and SemiBin(pretrain) with single-sample binning and VAMB and SemiBin with multi-sample binning in the human gut, dog gut and ocean datasets. Based on results from CheckM, SemiBin(pretrain) reconstruncted 55.7%, 94.6% and 49.7% more high-quality bins than Metabat2 with single-sample binning and 44.7%, 25.0% and 41.0% more high-quality bins than VAMB with multi-sample binning in the human gut, dog gut and ocean datasets, respectively.

**Supplementary Fig 15.**
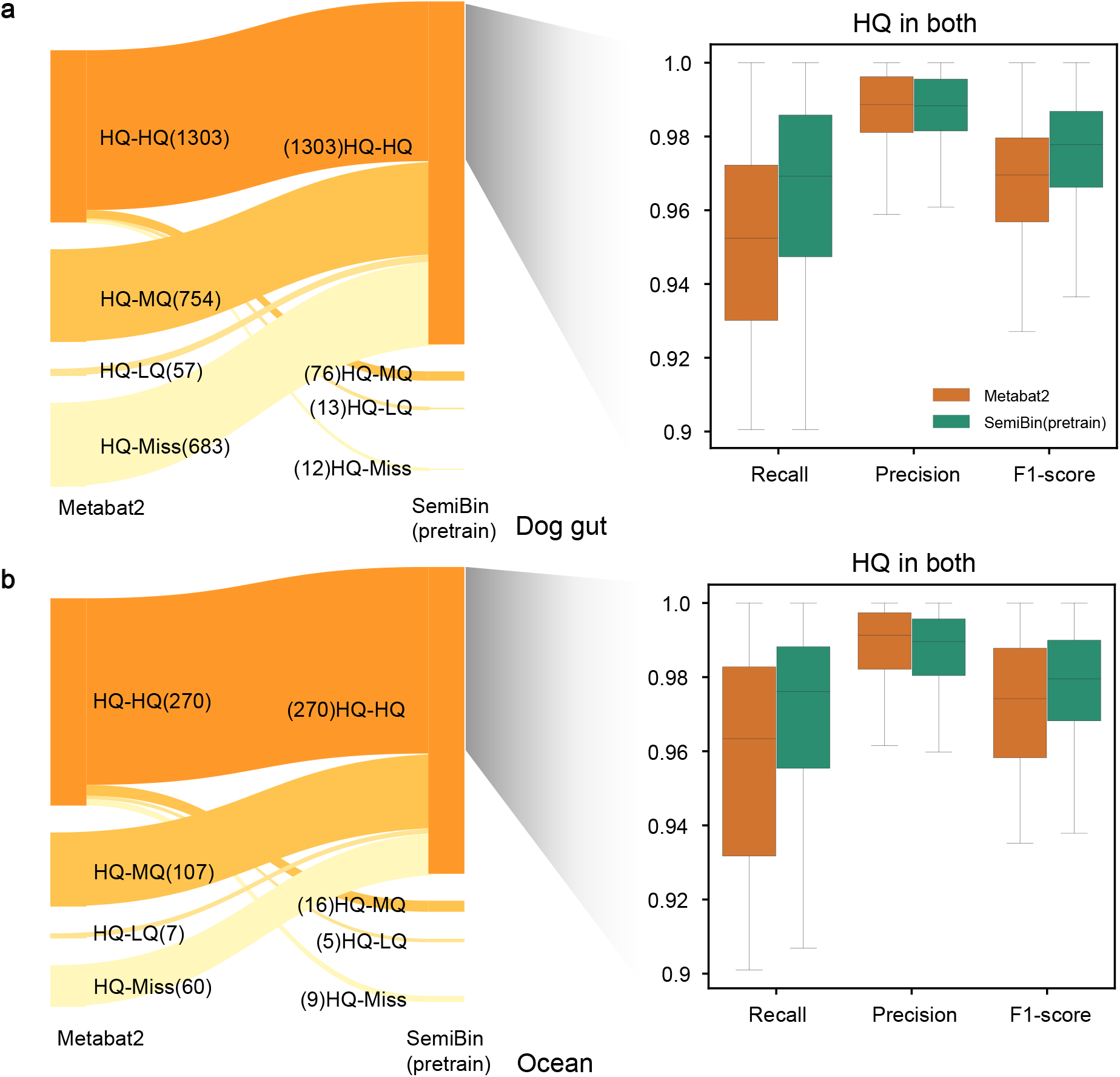
SemiBin(pretrain) reconstructed more and better high-quality bins compared to Metabat2 in dog gut (a) and ocean datasets (b). Results here are qualitatively similar to results of the human gut datasets (see Fig. 2c). In the dog gut and ocean datasets, within bins that are high-quality in both binners, SemiBin(pretrain) significantly achieved higher completeness (*P* = 7.338ů10^−110^; *P* = 5.266ů10^−20^) and F1 statistic (*P* = 1.107ů10^−110^; *P* = 4.331ů10^−15^), with slightly increase in contamination(*P* = 0.165 > 0.05; *P* = 3.953ů10^−05^; all *P*-values were computed from using Wilcoxon signed rank test, two-sided null hypothesis). (HQ-HQ: high-quality in both; HQ-MQ: high-quality in one and medium-quality in the other; HQ-LQ: high-quality in one and low-quality or worse in the other; HQ-Miss: high-quality in one and missed in the other)

**Supplementary Fig 16.**
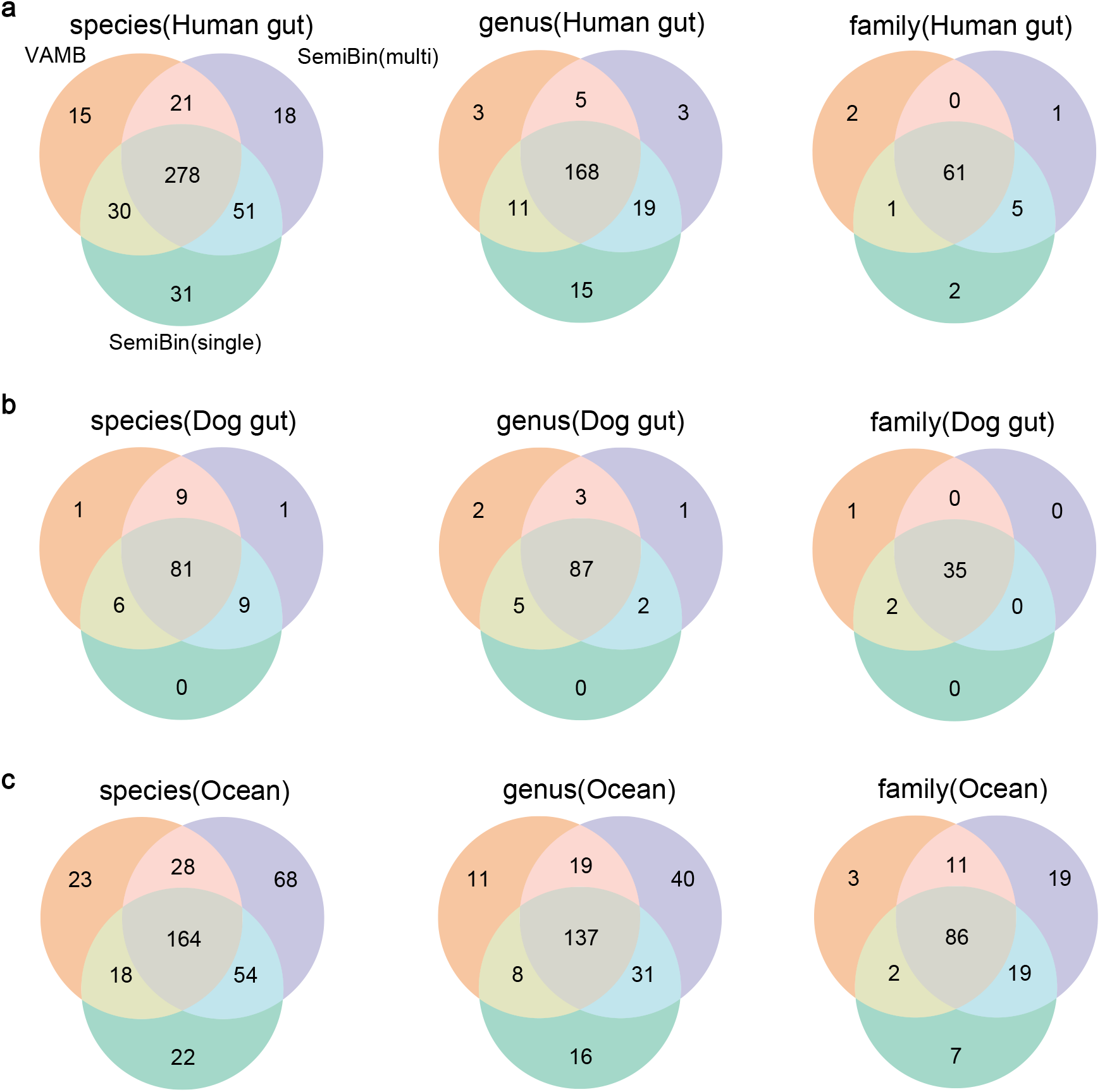
SemiBin reconstructed more high-quality distinct species, genera and families with multi-sample binning. We compared the the number of high-quality distinct species, genera and families of VAMB with multi-sample binning, SemiBin(multi) (SemiBin with multi-sample binning) and SemiBin(single) (SemiBin with single-sample binning) for **a,** human gut; **b,** dog gut; and **c,** ocean dataset.

**Supplementary Fig 17.**
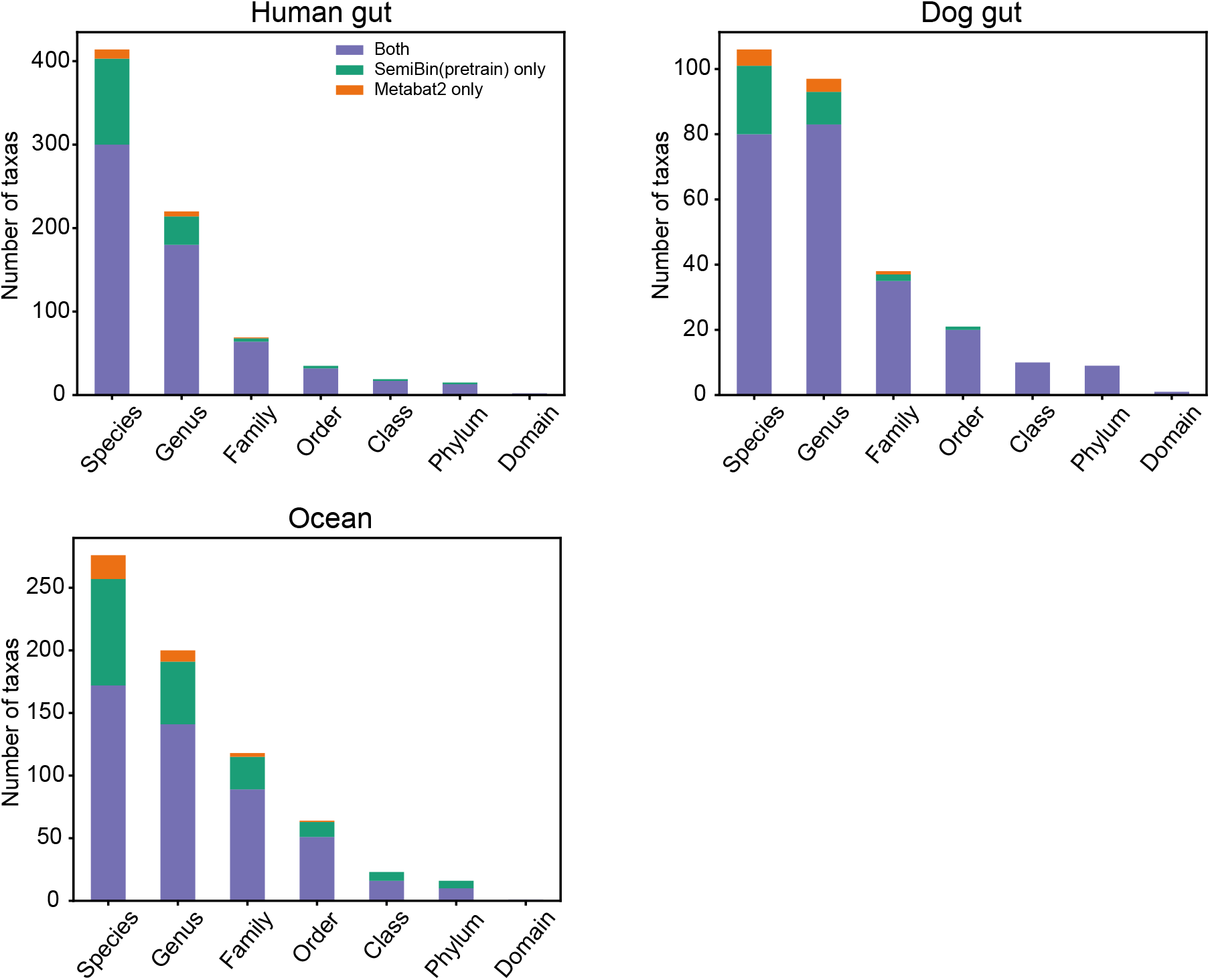
SemiBin(pretrain) had a larger taxonomic diversity at all levels compared to Metabat2 with single-sample binning in real datasets. We annotated the high-quality bins from SemiBin(pretrain) and Metabat2 with GTDB-TK. Shown is the number of distict taxa in the species, genus, family, order, class, phylum and domain level.

**Supplementary Fig 18.**
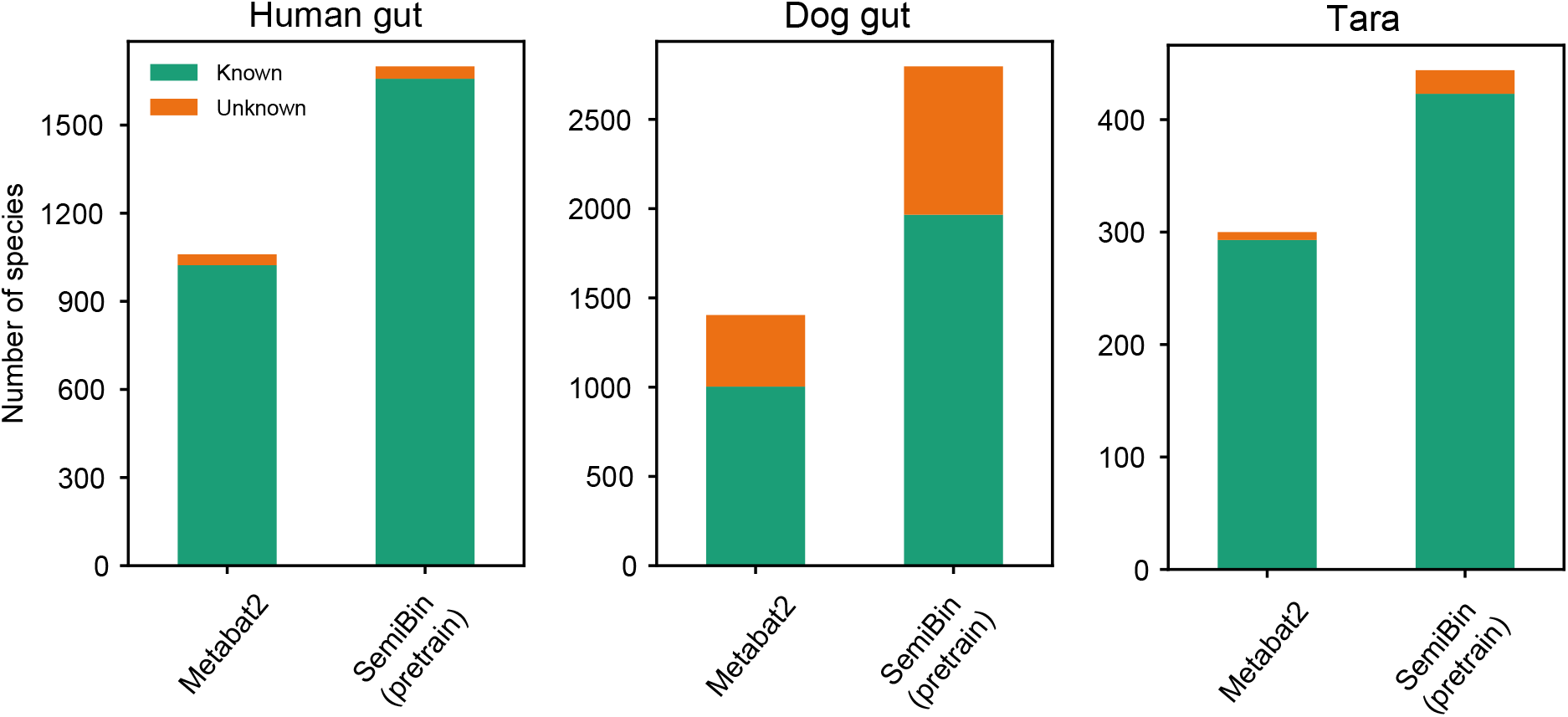
SemiBin(pretrain) reconstructed more known and unknown species. We annotated the high-quality bins from SemiBin(pretrain) and Metabat2 with GTDB-TK. Shown is the number of known and unknown species.

**Supplementary Fig 19.**
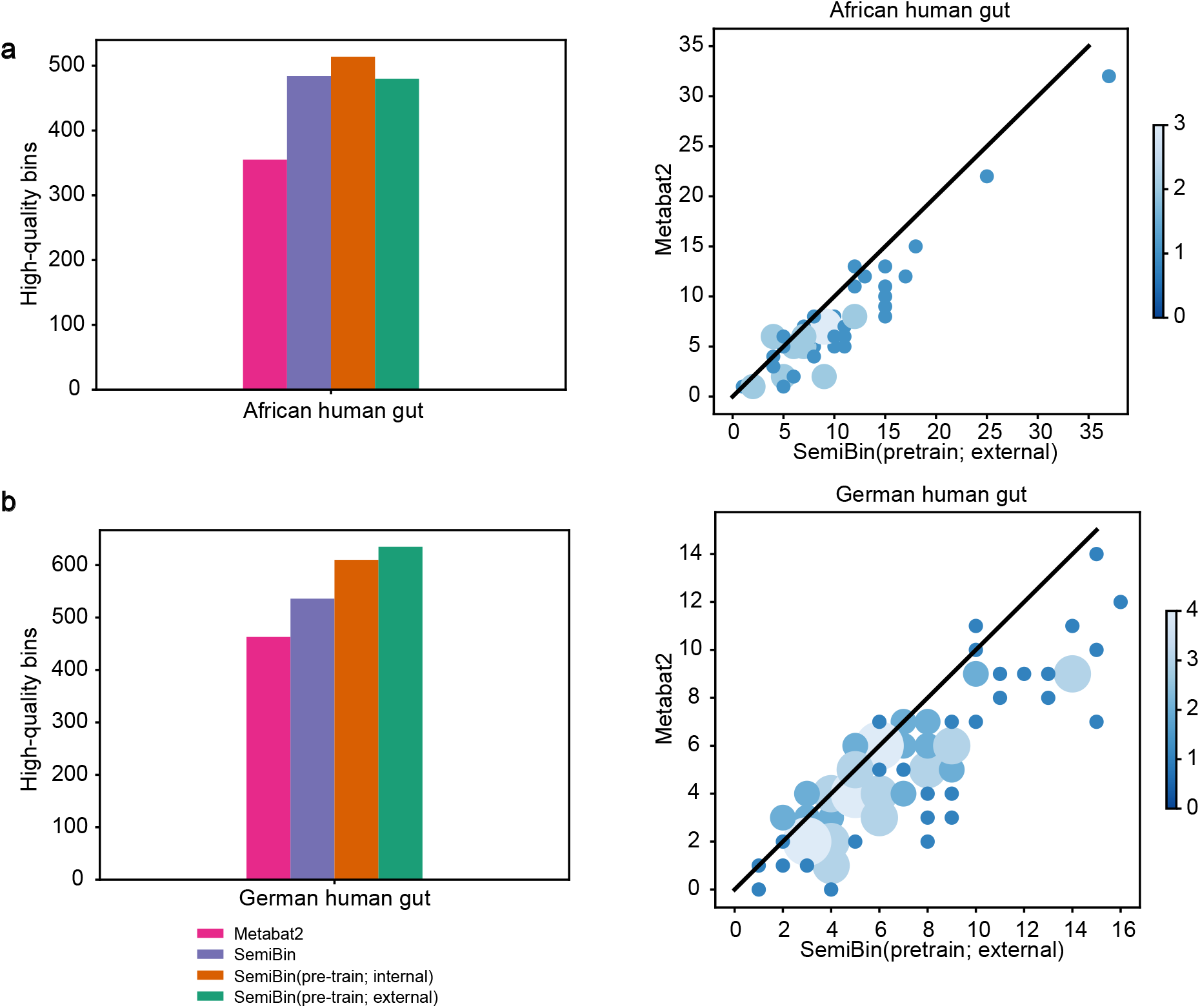
SemiBin with the pretrained model outperformed Metabat2 on two hold-out human gut datasets from African and German populations. We transferred the pretrained model from the human gut dataset to two hold-out human gut datasets from African and German populations. We also benchmarked Metabat2, SemiBin and SemiBin with the pretrained model that trained from the hold-out datasets (using 20 samples). (SemiBin(pretrain; internal): SemiBin with the pretrained model from the hold-out human gut datasets; SemiBin(pretrain; external): SemiBin with the pretrained model from the human gut datasets used in Fig. 2b. **a,** African human gut; **b,** German human gut.

**Supplementary Fig 20.**
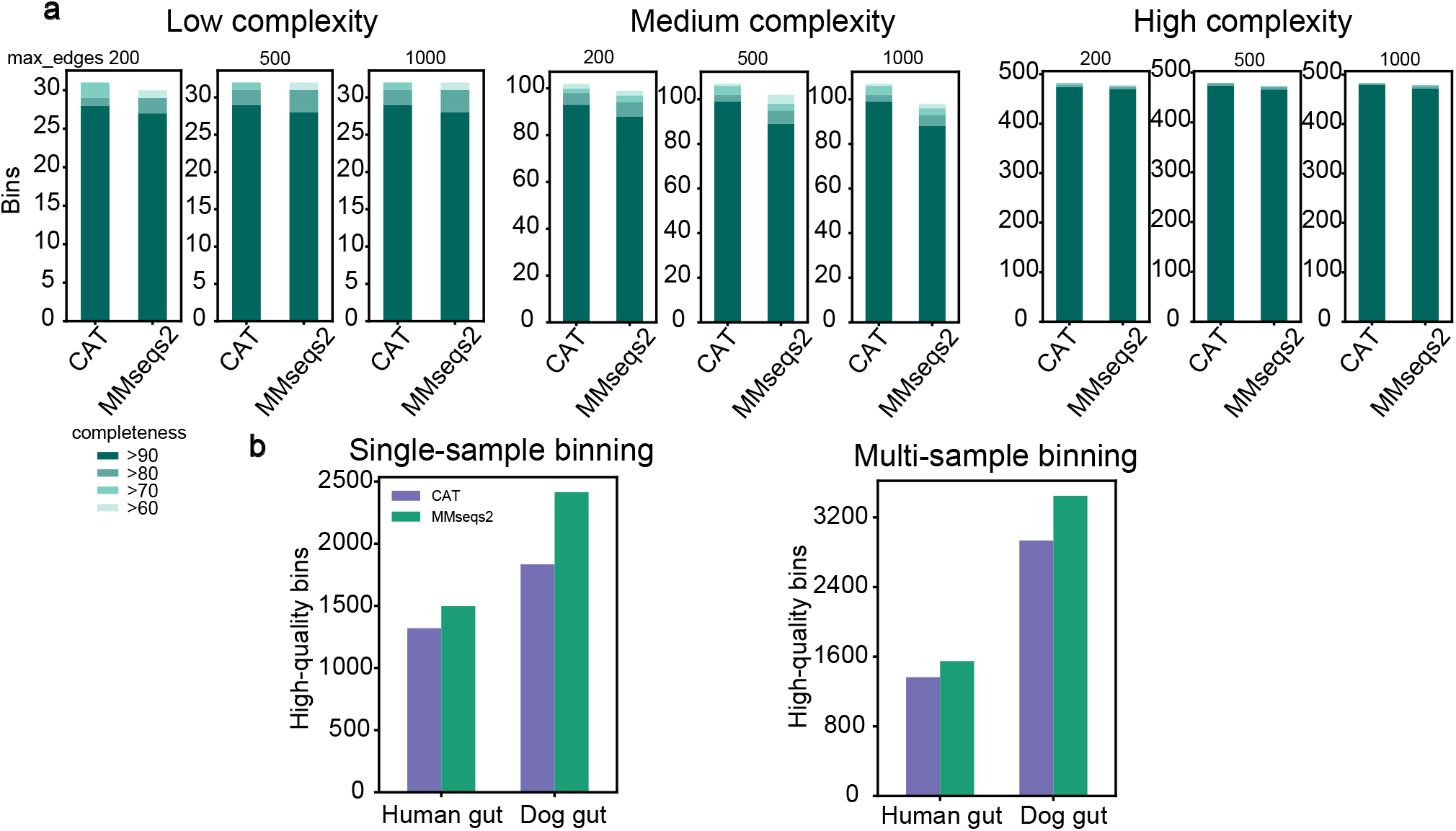
SemiBin with a better reference genome got better binning results. We compared the results of SemiBin with contig annotations by CAT(using NCBI reference genome) and MMseqs2(using GTDB reference genome) on **a,** CAMI I datasets and **b,** human gut and dog gut real datasets.

**Supplementary Fig 21.**
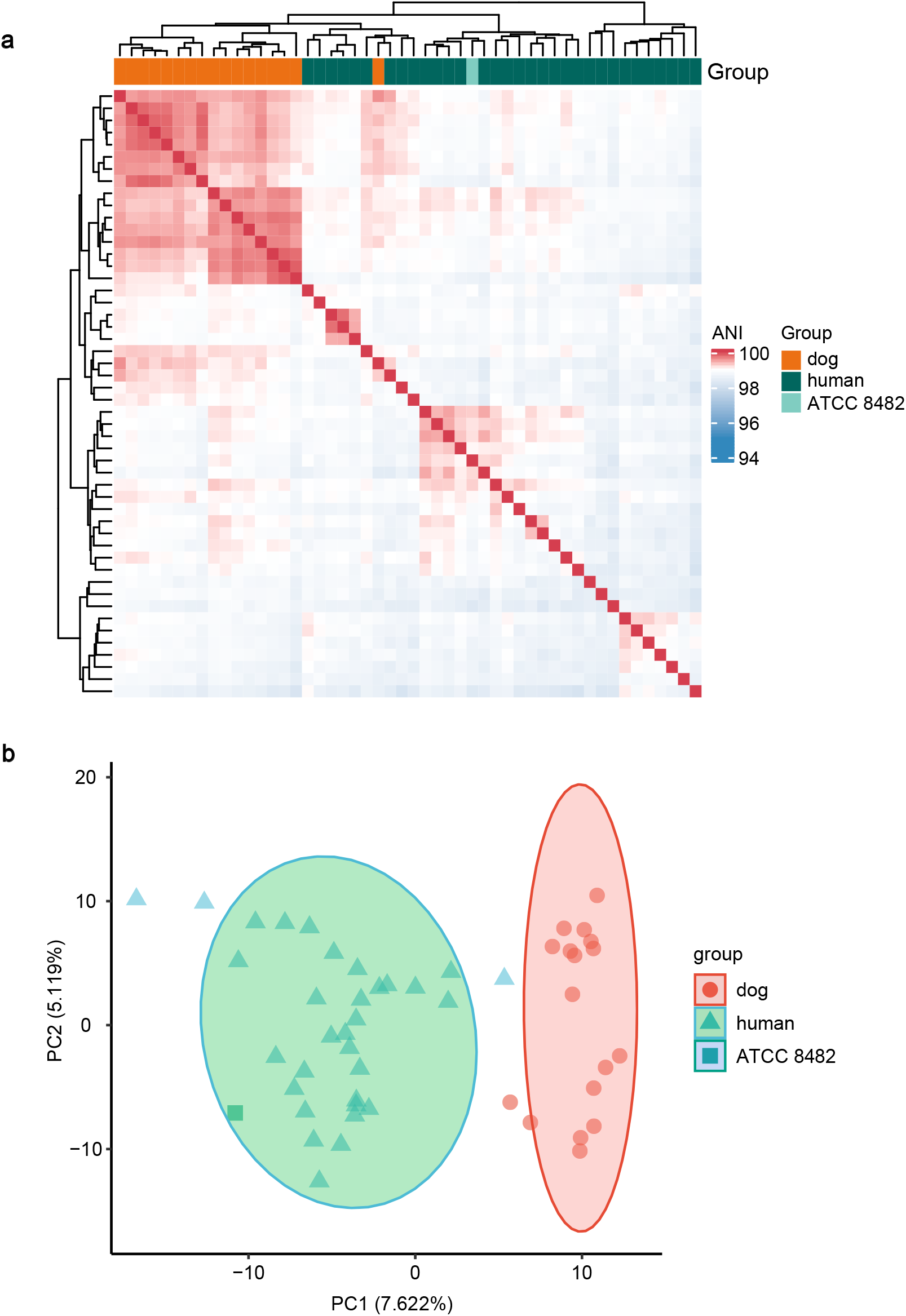
*B. vulgatus* bins from the dog gut clustered separately from those of the human gut. **a,**Heatmap representing the average nucleotide identity (ANI) similarity of the 49 strains of *B. vulgatus* studied and a type strain *B. vulgatus* ATCC 8482. Color scheme varies from high ANI similarity(red) to low ANI similarity(blue) of the strains analyzed. **b,** Principal component analysis (PCA) based on whole presence/absence genes.

**Supplementary Fig 22.**
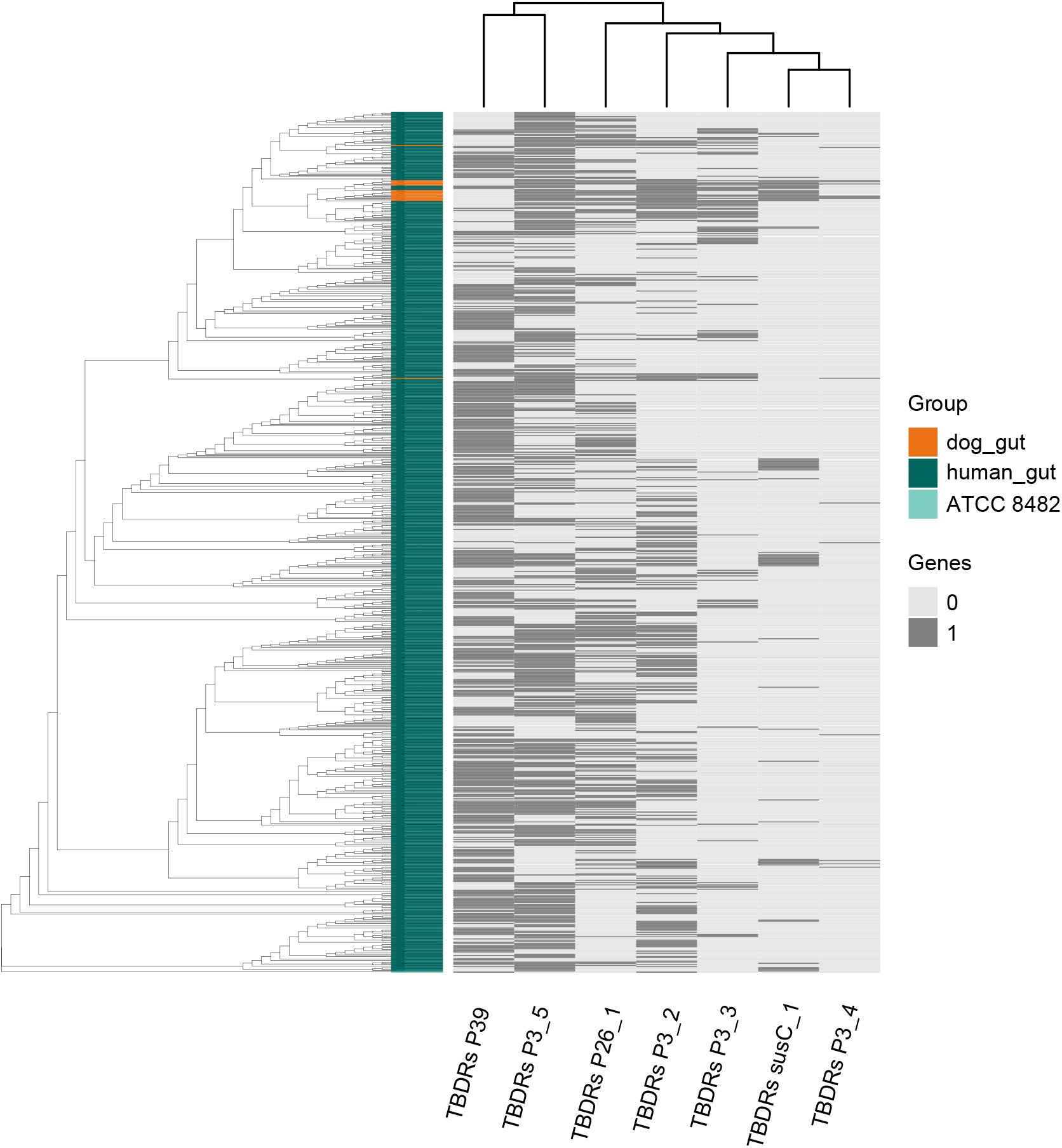
The separation of *B. vulgatus* strains in human gut and dog gut could also be found with GMGC as a external validation. We used 762 high-quality strains of *B. vulgatus* from the human gut in GMGC as the external validation and a type strain *B. vulgatus* ATCC 8482. Shown is the maximum-likelihood phylogenetic trees based on core genes and the heatmap showing the presence or absence of the genes encoding TonB-dependent receptors (TBDR).

**Supplementary Table 1.** Overview of the datasets used in the benchmarking.

|  | Dataset | #Samples | #Genomes | Binning mode |
| --- | --- | --- | --- | --- |
| CAMI I | Low complexity | 1 | 40 | Single-sample |
|  | Medium complexity | 2 | 132 | Co-assembly |
|  | High complexity | 5 | 596 | Co-assembly |
| CAMI II | Oral | 10 | 799 | Multi-sample |
|  | Skin | 10 | 610 | Multi-sample |
| Real datasets | Dog gut | 129 | Unknown | Single/Multi-sample |
|  | Human gut | 82 | Unknown | Single/Multi-sample |
|  | Tara | 109 | Unknown | Single/Multi-sample |

**Supplementary Table 2.** Accuracy of must-link and cannot-link constrains generated from contig annotations by MMseqs2 with GTDB reference genomes of simulated datasets. We used MMseqs2 with GTDB reference genome to annotate contigs of simulated datasets from CAMI I and CAMI II to test the results of the annotation. *Must-link Acc:* the accuracy of must-link constrains. *Cannot-link Acc:* the accuracy of cannot-link constrains. *#genomes:* number of genomes in the environment. *#Must-link covered genomes:* number of genomes that were covered by the accurate must-link constrains.

| Dataset | #must-link | Must-link Acc | #cannot-link | Cannot-link Acc | #genomes | #Must-link covered genomes |
| --- | --- | --- | --- | --- | --- | --- |
| Low | 2084477 | 0.980 | 12547725 | 0.988 | 40 | 31 |
| Medium | 3662242 | 0.992 | 76815282 | 0.986 | 132 | 92 |
| High | 206560 | 0.722 | 39533277 | 0.996 | 596 | 388 |
| Skin_sample_1 | 2346244 | 0.250 | 58684337 | 0.994 | 90 | 66 |
| Skin_sample_13 | 430244 | 0.170 | 3224103 | 0.982 | 140 | 35 |
| Skin_sample_14 | 580719 | 1.000 | 310471 | 0.887 | 58 | 6 |
| Skin_sample_15 | 2286104 | 0.125 | 14402383 | 0.988 | 72 | 39 |
| Skin_sample_16 | 220879 | 0.748 | 122596 | 0.537 | 41 | 5 |
| Skin_sample_17 | 806282 | 0.537 | 28461743 | 0.995 | 89 | 53 |
| Skin_sample_18 | 21817 | 0.646 | 191807 | 0.967 | 51 | 21 |
| Skin_sample_19 | 3432141 | 0.334 | 271153441 | 0.998 | 241 | 154 |
| Skin_sample_20 | 1843334 | 0.133 | 37709100 | 0.996 | 163 | 106 |
| Skin_sample_28 | 109276 | 0.999 | 213438 | 0.895 | 25 | 3 |
| Oral_sample_6 | 148247 | 0.971 | 4538540 | 0.974 | 89 | 46 |
| Oral_sample_7 | 302528 | 0.285 | 11557221 | 0.995 | 179 | 109 |
| Oral_sample_8 | 475445 | 0.294 | 20965330 | 0.996 | 163 | 96 |
| Oral_sample_13 | 491869 | 0.391 | 27319826 | 0.996 | 164 | 93 |
| Oral_sample_14 | 696582 | 0.548 | 39416022 | 0.997 | 141 | 88 |
| Oral_sample_15 | 543180 | 0.529 | 28024955 | 0.997 | 165 | 100 |
| Oral_sample_16 | 570645 | 0.573 | 25082966 | 0.994 | 320 | 85 |
| Oral_sample_17 | 349459 | 0.294 | 16970935 | 0.995 | 143 | 77 |
| Oral_sample_18 | 570501 | 0.678 | 33019031 | 0.997 | 150 | 82 |
| Oral_sample_19 | 612717 | 0.369 | 31035653 | 0.995 | 274 | 128 |

**Supplementary Table 3.** Binning tools used in recent large-scale metagenomy studies.

|  | #samples | Habitat(s) | Binning tool |
| --- | --- | --- | --- |
| Pasoli et al. (Pasolli et al., 2019) | 9,428 | Human associated | Metabat2 |
| Coelho et al. | 13,174 | Global microbiome | Metabat2 |
| Almeida et al. (Almeida et al., 2019) | 11,850 | Human gut | Metabat2 |
| Nayfach et al. (Nayfach et al., 2019) | 3,810 | Human gut | DAS Tool(Maxbin2;Metabat2;CONCOCT) |
| Nayfach et al. (Nayfach et al., 2020) | > 10,000 | Global microbiome | Metabat |
| Schulz et al. (Schulz et al., 2020) | 8,535 | Global virome | Metabat2 |
| Asnicar et al. (Asnicar et al., 2021) | 1,203 | Human gut | Metabat2 |

**Supplementary Table 4.** Running time and memory usage of SemiBin with different binning mode. For every environment, we randomly chose 10 samples to evaluate the running time and memory usage of SemiBin. Shown are the average running time and memory usage of ten samples except generating data with multi-sample binning, which generated data for all samples at the same time. For CPU machine, we used AWS g4ad.4xlarge machine, and for GPU machine, we used Tesla T4. GPU: Graphical Processing Unit, CPU: Central Processing Unit.

|  |  | Single-sample |  |  | Single-sample(pretrain) |  |  | Multi-sample |  |  |
| --- | --- | --- | --- | --- | --- | --- | --- | --- | --- | --- |
|  |  | Human gut | Dog gut | Ocean | Human gut | Dog gut | Ocean | Human gut | Dog gut | Ocean |
| Generating data | Time(min) | 6 | 10 | 16 | 6 | 10 | 16 | 29 | 36 | 36 |
|  | Memory(MB) | 1,229 | 1,119 | 2,894 | 1,229 | 1,119 | 2,894 | 1,182 | 1,147 | 1,219 |
| Generating cannot-link | Time(min) | 88 | 81 | 114 | n/a | n/a | n/a | 88 | 81 | 114 |
|  | Memory(MB) | 37,740 | 37,218 | 41,585 | n/a | n/a | n/a | 37,740 | 37,218 | 41,585 |
| Training(CPU) | Time(min) | 181 | 209 | 222 | n/a | n/a | n/a | 184 | 210 | 222 |
|  | Memory(MB) | 2,099 | 2,215 | 2,744 | n/a | n/a | n/a | 2,101 | 2,216 | 2,754 |
| Training(GPU) | Time(min) | 34 | 36 | 45 | n/a | n/a | n/a | 34 | 36 | 45 |
|  | Memory(MB) | 4,095 | 4,216 | 4,748 | n/a | n/a | n/a | 4,097 | 4,218 | 4,755 |
| Binning | Time(min) | 5 | 6 | 8 | 5 | 6 | 8 | 9 | 10 | 16 |
|  | Memory(MB) | 8,344 | 7,949 | 18,002 | 8,344 | 7,949 | 18,002 | 4,798 | 4,757 | 9,452 |

